# Mitochondrial pyruvate metabolism regulates the activation of quiescent adult neural stem cells

**DOI:** 10.1101/2022.05.31.494137

**Authors:** Francesco Petrelli, Valentina Scandella, Sylvie Montessuit, Nicola Zamboni, Jean-Claude Martinou, Marlen Knobloch

## Abstract

Cellular metabolism is important for adult neural stem/progenitor cell (NSPC) behavior. However, its role in the transition from quiescence to proliferation is not fully understood. We here show that the mitochondrial pyruvate carrier (MPC) plays a crucial and unexpected part in this process. MPC transports pyruvate into mitochondria, linking cytosolic glycolysis to mitochondrial tricarboxylic acid cycle (TCA) and oxidative phosphorylation (OXPHOS). Despite its metabolic key function, the role of MPC in NSPCs has not been addressed. We show that quiescent NSPCs have an active mitochondrial metabolism and express high levels of MPC. Pharmacological MPC inhibition increases aspartate and triggers NSPC activation. Furthermore, genetic MPC-ablation *in vivo* also activates NSPCs, which differentiate into mature neurons, leading to overall increased hippocampal neurogenesis in adult and aged mice. These findings highlight the importance of metabolism for NSPC regulation and identify a novel pathway through which mitochondrial pyruvate import controls NSPC quiescence and activation.

**Highlights:**

• Quiescent NSPCs have high levels of MPC and an active mitochondrial network
• The import of pyruvate into mitochondria is necessary to maintain quiescence of NSPCs
• MPC inhibition increases intracellular aspartate levels and triggers the activation of quiescent NSPCs
• MPC-knockout NSPCs generate mature newborn neurons, leading to overall increased neurogenesis in adult and advanced age mice

**Graphical abstract:** 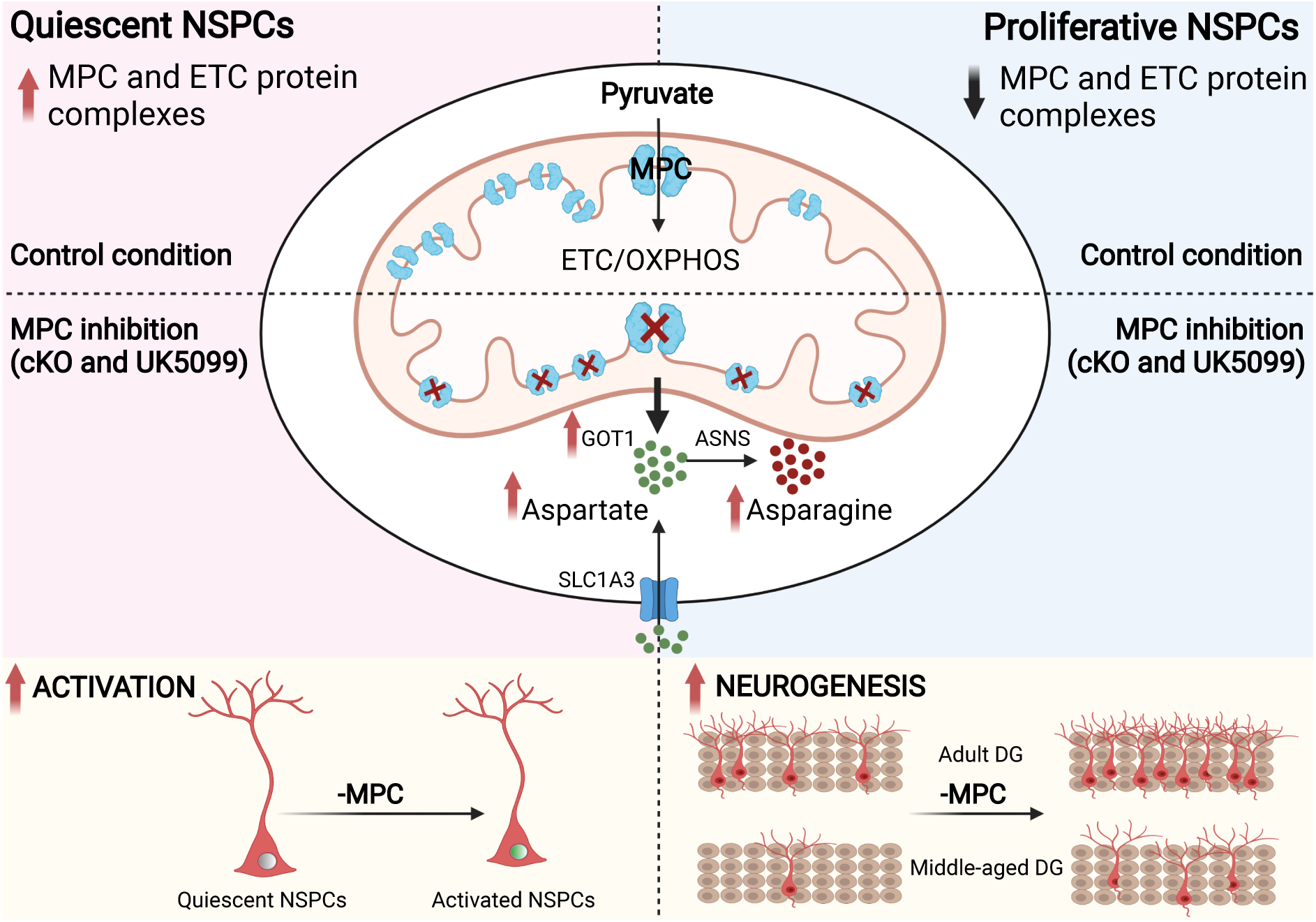

## Introduction

Stem cells must maintain a tight balance between quiescence, proliferation and differentiation to sustain lifelong tissue regeneration and maintenance. This is also the case for adult NSPCs, which form newborn neurons throughout life (Denoth-Lippuner and Jessberger, 2021). NSPCs are primarily quiescent in adulthood, but can proliferate upon intrinsic and extrinsic stimuli (Urbán and Cheung, 2021). Although NSPC activation is critical for proper neurogenesis, the underlying mechanisms are still not fully understood.

Cellular metabolism has been shown to determine the activity state of stem cells (Cavallucci et al., 2016; Ly et al., 2020) and metabolic features appear similar among different tissue-specific adult stem cells. Quiescence is considered as a state in which stem cells mainly use glycolysis to sustain their energy demand. Along with the differentiation process, cells shift from a more glycolytic profile to a more oxidative metabolism (Shapira and Christofk, 2020). Such a metabolic shift seems also important for neural stem/progenitor cells (NSPCs). Indeed, single cell RNA sequencing (scRNA-Seq) studies found decreased expression of glycolytic genes and an upregulated expression of genes involved in oxidative phosphorylation (OXPHOS) at early stages of fate transition in adult NSPCs (Beckervordersandforth et al., 2017; Shin et al., 2015). These findings are supported by metabolic analyses of NSPCs under differentiation *in vitro* (Zheng et al., 2016). However, this classical view of metabolic quiescence has been challenged with recent findings showing that quiescent NSPCs have high levels of mitochondrial fatty acid beta-oxidation (FAO) and express many proteins involved in diverse aspects of mitochondrial metabolism (Knobloch et al., 2017; Wani et al., 2022). Furthermore, mitochondria are abundant in NSPCs and their dynamics affects self-renewal and fate choice (Iwata et al., 2020; Khacho et al., 2016). Thus, mitochondrial metabolism might play a more important role for NSPC quiescence than previously anticipated.

A shift from a glycolytic to more oxidative metabolism very often requires a redirection of pyruvate, the end product of glycolysis, from lactate production towards mitochondrial oxidation in the TCA cycle. The mitochondrial pyruvate carrier (MPC), a heterodimer of MPC1 and MPC2, is required for this transport (Bricker et al., 2012; Herzig et al., 2012; Vanderperre et al., 2016). Absence of one of the two proteins leads to a loss of pyruvate transport and has a profound impact on the metabolic state of the cells (Vanderperre et al., 2015; Zangari et al., 2020). Despite its key role in linking glycolysis and mitochondrial metabolism, it remains unknown whether MPC plays a regulatory role in NSPC behavior.

Therefore, we here used pharmacological inhibition and genetic deletion of MPC to assess whether a disruption of pyruvate import into the mitochondria would affect NSPC maintenance, activation, and differentiation. Surprisingly, we found that quiescent NSPCs express high levels of MPC and require pyruvate import into the mitochondria for the maintenance of quiescence. Inhibition of MPC triggers their activation by increasing the intracellular pool of aspartate, despite a significant decrease of TCA cycle intermediates. Furthermore, MPC-knockout NSPCs are able to differentiate into mature neurons, indicating a high metabolic flexibility, which allows these cells to adapt their metabolism according to substrate availability. We further show that this increased activation and undisturbed differentiation of MPC-knockout NSPCs leads to an overall increase in neurogenesis in adult and middle-aged mice.

## Results

### MPC is dynamically regulated with activity state and its transport function is required for NSPC quiescence

In order to determine the role of MPC in NSPCs, we first analyzed the expression of MPC1 in existing RNA-Seq databases. We found that MPC1 is expressed in NSPCs and in other cell types in the dentate gyrus (DG) of adult mice (Fig. 1A) (Hochgerner et al., 2018). Available images from dentate gyrus sections of MPC1-GFP reporter mice (www.gensat.org) confirmed the expression of MPC1 at protein level in various cell types of the neurogenic niche, including NSPCs (Suppl. Fig 1A).

**Figure 1.**
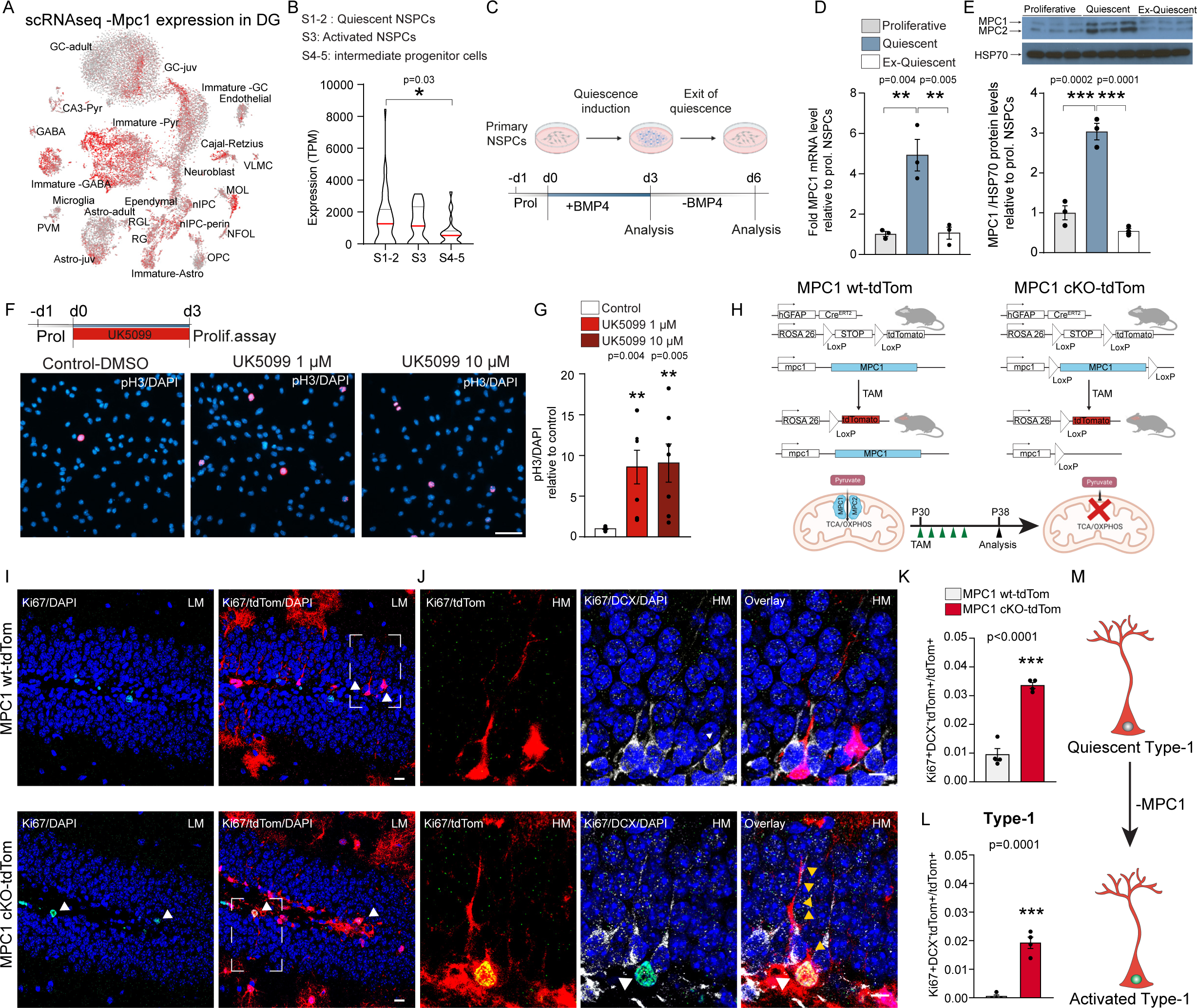
MPC is necessary to maintain the quiescence in NSPCs. A. t-SNE visualization of Mpc1 expression in the DG of adult mice. Single cell RNAseq data were queried using an online resource (linnarsonnlab.org/dentate, Hochgerner et al. Nat. Neuro. 2018) B. Violin plots of Mpc1 expression in quiescent (S1-2), more activated NSPCs (S3) and intermediate progenitor cells (S4-5). Data queried from Shin et al. Cell Stem Cell 2015. Red line represents mean. *p<0.05, **p<0.01, ***p< 0.001 (one-way ANOVA followed by post-hoc test) C. Illustration of the artificial reversible system to induce NSPC quiescence *in vitro*. D. Relative mRNA expression of *Mpc1* in proliferative (grey), quiescent (light blue) and ex-quiescent (white) NSPCs. *Mpc1* is expressed as fold change to proliferative NSPCs, bars represent mean ± SEM. **p<0.01, n=3 biological replicates (one-way ANOVA followed by post-hoc test). E. Representative western blots of MPC1 (12 kDa), MPC2 (14 kDa) and HSP70 (70KDa) in proliferative (grey), quiescent (light blue) and ex-quiescent (grey) NSCs. MPC1 protein levels were normalized to HSP70 levels and expressed as fold change to proliferative NSPCs. Bars represent mean ± SEM. ***p<0.0001, n=3 biological replicates (one-way ANOVA followed by post-hoc test). F. Top: Experimental outline of UK5099 treatment during quiescence induction. Bottom: Representative images of pH3 (red) and DAPI (blue) immunostaining in quiescent NSPCs treated with UK5099. Scale bar: 50 μm. G. Quantification of pH3+ in DMSO (white) or UK5099 (light and dark red) treated quiescent NSPCs. Data represent mean ± SEM. *p<0.05. n=6 CS from 3 independent experiments (one-way ANOVA followed by post-hoc test) H. Scheme showing the generation of MPC1 cKO-tdTom and MPC1 wt-tdTom mice. I. Representative low magnification (LM) confocal images of TAM-induced td-Tomato recombination (red), Ki67 (green) and DAPI (blue) immunostaining at P38 in DG of control MPC1 wt-tdTom (grey) and MPC1 cKO-tdTom (red) mice. Scale bars: 10 μm J. Representative high magnification (HM) confocal images of TAM-induced td-Tomato recombination (red), Ki67 (green), doublecortin (DCX, white) and DAPI (blue) immunostaining at P38 in the DG of control MPC1 wt-tdTom (grey) and MPC1 cKO-tdTom (red) mice. White arrowheads point show the colocalization of Ki67 and tdTom in a single confocal plane. Yellow arrowheads show type-1 cell processes. Scale bars: 10 μm. K. Top: Experimental scheme. TAM injection at P30 (green arrow) for 5 consecutive days and data analysis at P38 (black arrow). Below: Quantification of Ki67 positive (Ki67+) tdTom+ cells over total number of tdTom+ cells in the DG of and control MPC1 wt-tdTom (grey) an MPC1 cKO-tdTom (red) mice. Data are shown as mean value ± SEM. **p<0.01, n=4 mice per group (unpaired Student’s t-test). L. Quantification of type-1 Ki67+ DCX-tdTom+ cells over the total number of tdTom+ cells in MPC1 wt-tdTom (grey) and MPC1 cKO-tdTom (red) mice. Data are shown as mean ± SEM. ***p<0.001, n=4 mice per group (unpaired Student’s t-test). M. Illustration shows type-1 cell activation following MPC1 deletion

Surprisingly, when we looked at scRNA-Seq data comprising different activity states of NSPCs *in vivo* (Shin et al., 2015) we found that MPC1 expression was highest in the more quiescent NSPC populations and decreased with lineage progression (Fig 1B and Suppl Fig 1B), suggesting that the import of pyruvate into the mitochondria might play a fundamental role in the regulation of the quiescent state in NSPCs.

To further assess whether MPC1 expression was dependent on the activity state of NSPCs, we used cultured adult hippocampal NSPCs and applied an established *in vitro* system, which mimics quiescence in a reversible way (Fig. 1C) (Knobloch et al., 2017; Martynoga et al., 2013; Mira et al., 2010). Supporting the scRNA-Seq analysis, we found that the levels of MPC1 mRNA and protein were significantly higher in quiescent NSPCs compared to active/proliferating NSPCs (Fig. 1D and E). MPC2 followed the same pattern (Fig. 1E and Suppl. Fig 1C). When the artificial quiescence state of NSPCs was reverted back to a proliferative state, MPC1 and MPC2 levels were significantly reduced to the levels of proliferative NSPCs (Fig. 1D and E and Suppl Fig C), indicating that MPC levels are dynamically regulated with activity state. OXPHOS complexes followed the same pattern, with increased protein levels in quiescent NSPCs (Suppl. Fig. 1D and E). Furthermore, morphometric analysis of mitochondria revealed that quiescent NSPCs contained primarily fused and elongated mitochondria, which are indicative of highly active OXPHOS (Alirol and Martinou, 2006; Giacomello et al., 2020), whereas proliferative NSPCs had more fragmented mitochondria (Suppl. Fig. 1F and G). Taken together, these findings show that quiescent NSPCs have high MPC expression and an elongated mitochondrial network.

Given the high MPC levels, we next tested whether inhibiting MPC would impact on the quiescence of these cells. We used the specific MPC inhibitor UK5099 (Halestrap, 1975) either during quiescence induction (Fig 1F and G) or after established quiescence (Suppl Fig 1H and I). Strikingly, UK5099 significantly increased the number of cycling and dividing Ki67 and phospho-Histone 3 (pH3) positive NSPCs, suggesting that MPC activity is indeed required for maintenance of quiescence and that its blockage leads to a NSPC activation *in vitro*.

### MPC1 deletion in NSPCs *in vivo* leads to increased numbers of progeny and triggers NSPC proliferation

We next addressed whether MPC is important for the maintenance of quiescent NSPCs *in vivo*. To delete MPC1 in adult NSPCs, we crossed MPC1 floxed mice (MPC1 fl/fl, (Gray et al., 2015)) with a tamoxifen-inducible GFAP-Cre recombinase line (hGFAP-CreERT^2^, Hirrlinger et al., 2006). To visualize the recombined cells and their progeny, we additionally crossed these inducible MPC1 knockout mice with an inducible tdTomato lineage tracing line (tdTom, Madisen et al., 2010), resulting in triple transgenic mice hereafter called MPC1 cKO-tdTom mice (Fig. 1H). Control mice were generated the same way, but carried wildtype alleles of the MPC1 gene (MPC1 wt-tdTom mice). Cre-expression was induced by administration of tamoxifen at postnatal day 30 (P30), leading to deletion of MPC1 in GFAP-expressing cells from cKO-tdTom mice, including NSPCs (Fig 1K). To determine the effect of MPC1 deletion on the activation of NSPCs *in vivo*, we analyzed the number of tdTomato positive cells at P38 (Fig 1I).

While the total number of tdTom+ cells was not increased at P38, i.e. eight days after the first tamoxifen injections (Suppl Fig1J and K), the proportion of activated/proliferative tdTom+ cells was significantly increased, as revealed by the cell cycle marker protein Ki67 (Fig 1J and K). As several subtypes of NSPCs in the DG can proliferate, we categorized the Ki67+/ tdTom+ cells according to the previously described nomenclature (Kempermann et al., 2004) into type-1 cells (radial glia-like cells) and type-2a cells (no radial process, more proliferative progenitors). Type-2a cells were further distinguished from committed neuroblasts by the absence of the protein doublecortin (DCX), an immature neuronal marker. Whereas the number of Ki67+ tdTom+ type-2a cells was comparable between control and MPC1 cKO-tdTom mice (Suppl. Fig 1L), the number of Ki67+tdTom+ cells type-1 cells was significantly increased in the KO mice (Fig. 1J and L), indicating a substantial activation of quiescent NSPCs upon MPC1 deletion (Fig 1M).

Taken together, these data show that MPC1 deletion in NSPCs leads to an increased activation and proliferation of NSPCs, resulting in higher numbers of progeny generated. These findings are in line with the increased activation of quiescent NSPCs seen *in vitro* upon MPC1 blockage (Fig. 1F and G).

### MPC1 inhibition activates quiescent NSPCs by increasing the intracellular pool of aspartate

We next assessed the metabolic profile of quiescent NSPCs by using Seahorse technology. A fuel flexibility test showed that the oxygen consumption rate (OCR) in quiescent NSPCs depends mainly on glucose. However, glutamine and fatty acids also significantly contributed as oxidative fuel sources (Suppl Fig 2A), suggesting that quiescent NSPCs are flexible in their use of substrates for energy and biosynthetic metabolism. OCR was markedly impaired after acute injection of UK5099 under basal conditions in quiescent NSPCs (Fig 2A) while the inhibition of FAO and glutaminolysis using etomoxir and BPTES respectively, decreased OCR to a lesser extent (Fig Suppl 2C). Acute inhibition with UK5099 also led to an increase in the extracellular acidification rate (ECAR) and lactate secretion in quiescent NSPCs (Fig. 2A and Suppl Fig 2B and D). Together these data suggest that quiescent NSPCs use pyruvate as the main substrate for their mitochondrial respiration, and that at least a part of the pyruvate produced by glycolysis is transported into the mitochondria rather than being converted to lactate.

**Figure 2:**
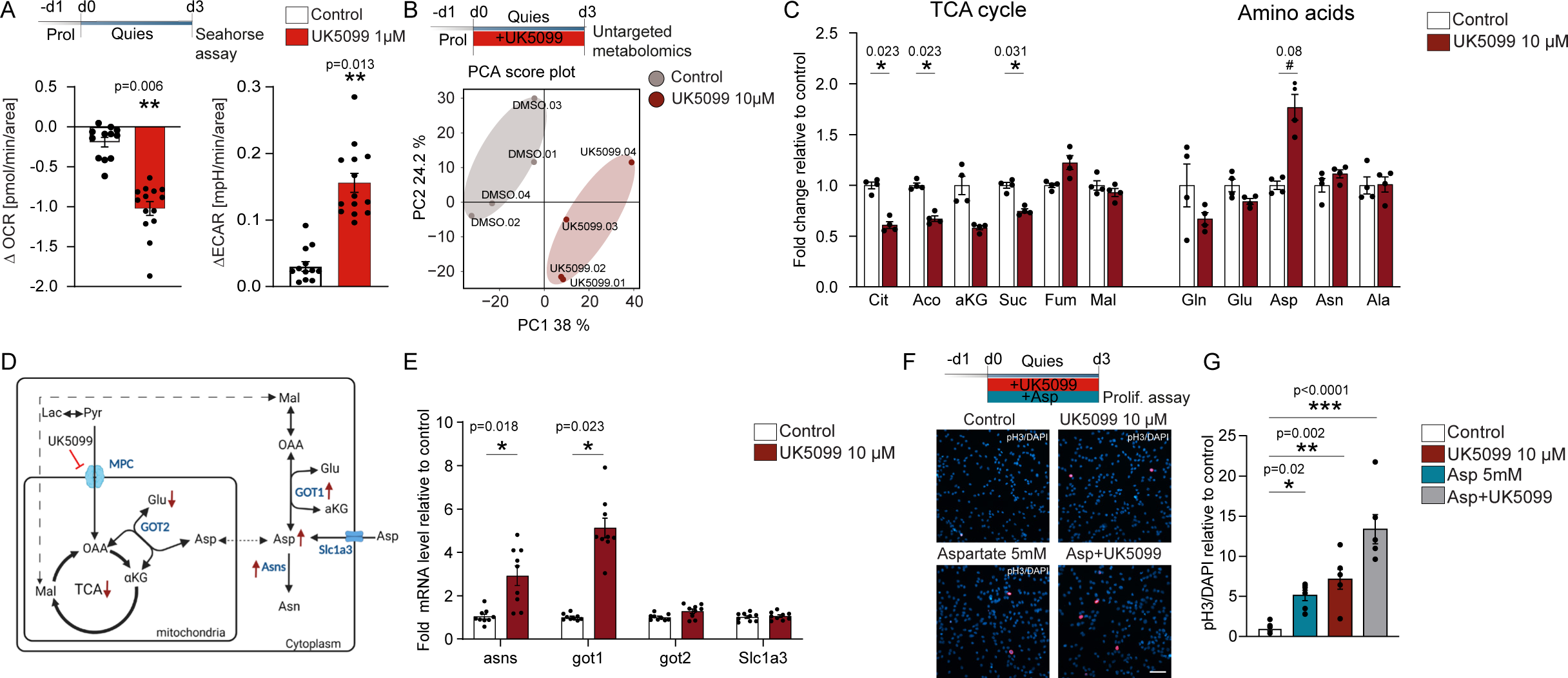
MPC1 inhibition activates quiescent NSPCs by increasing the intracellular pool of aspartate. A. Experimental outline of seahorse analysis in quiescence NSPCs. Left graph represents OCR changes after acute MPC inhibition with UK5099 1 μM. Right graph represents ECAR changes after acute MPC inhibition. Bars represent mean ± SEM.**p<0.01, n=10-15 wells from 3 independent experiments. Statistics were computed on averaged experiments (unpaired Student’s t-test). B. Top: Experimental outlines of untargeted metabolomics in quiescent NSPCs treated with UK5099. Bottom: Principal component analysis (PCA) of untargeted metabolomics from control and UK5099-treated quiescent NSPCs. Grey represents control group and red represents UK5099 treated cells. N= 4 biological replicates. C. Relative intensity of selected metabolites in control and UK5099 treated quiescent NSPCs. Bars represent mean ± SEM. * adjusted p-value <0.05, # adjusted p-value <0.1, N=4 biological replicates. Cit: Citrate, Aco:Aconitate, aKG: alpha-ketoglutarate, Suc: Succinate, Fum: Fumarate, Mal: Malate, Gln: Glutamine, Glu: Glutamate, Asp: Aspartate, Asn: Asparagine, Ala: Alanine. D. Scheme of the main metabolic pathways involved in the synthesis and utilization of aspartate. Red arrows indicate up and downregulated enzymes upon UK5099 treatment (see E). E. Relative mRNA expression of *Asns, Got1, Got2* and *Slc1a3* in control (white) and UK5099-treated (red) NSPCs. Genes are expressed as fold change to control NSPCs, bars represent mean ± SEM. * p<0.05, N=9 biological replicates from 3 independent experiments. Statistics were computed on averaged experiments (unpaired Student’s t-test). F. Top: Experimental outline of UK5099 and aspartate treatment during quiescence induction. Bottom: Representative images of pH3 (red) and DAPI (blue) immunostaining in quiescent NSPCs treated with UK5099, Aspartate and UK5099+Aspartate. Scale bar: 50 μm. G. Quantification of pH3+ in control (white), Aspartate (blue), UK5099 (red) and Aspartate+UK5099 (grey) treated quiescent NSPCs. Data represent mean ± SEM. *p<0.05, **p<0.01, ***p<0.005. n=6 CS from 2 independent experiments (one-way ANOVA followed by post-hoc test)

To investigate whether the block of mitochondrial pyruvate import in quiescent NSPCs rewires their metabolism, we next performed untargeted metabolomics on NSPCs cultured in the presence or absence of UK5099 during the induction of quiescence (Fig 2B). Principal component analyses revealed that samples clustered according to the treatment, suggesting that MPC inhibition drives profound metabolic changes in quiescent NSPCs (Fig 2B and Suppl Fig 2E). Pathway enrichment analysis showed that UK5099 significantly affected TCA cycle and amino acid metabolism (Suppl Fig 2F). While intermediates of glycolysis were not significantly changed (Suppl Fig 2G), TCA intermediates, in particular citrate and succinate levels, were clearly reduced compared to untreated cells. Of note, malate and fumarate were not significantly different (Fig 2C). Furthermore, MPC inhibition lead to a significant increase of intracellular levels of aspartate (Fig 2C). Overall, these measurements indicate an impairment in TCA activity and a rewiring of amino acid metabolism towards aspartate production (Fig 2C).

The intracellular pool of aspartate is maintained by the activity of two transaminases, the cytosolic enzyme glutamic-oxaloacetic transaminase 1 (GOT1) and the mitochondrial glutamic-oxaloacetic transaminase 2 (GOT2). GOT1 uses aspartate to generate glutamate and oxaloacetate, while GOT2 uses these two substates to synthesize α-ketoglutarate and aspartate (Fig 2D). In the absence of ETC activity, GOT1 reverts its reaction in order to produce aspartate (Birsoy et al., 2015). We thus measured the mRNA expression of GOT1 and GOT2. GOT1 transaminase was significantly increased in quiescent NSPCs treated with UK5099 (Fig 2E). In addition, asparagine synthase (Asns), which converts aspartate to asparagine was also increased (Fig 2E). These data suggest that increased expression of GOT transaminases upon MPC inhibition contributes to the increased aspartate levels.

Cells can also uptake aspartate through the glutamate-aspartate plasma membrane transporter Solute Carrier Family 1 Member 3 (SLC1A3, also known as GLAST) (Fig 2D). Quiescent NSPCs express already high levels of this transporter (Mori et al., 2006), and no further increase occurred upon UK5099 treatment (Fig 2E). Remarkably, exogenous aspartate or overexpression of SLC1A3 has been shown to be sufficient to support proliferation of cells lacking ETC activity (Birsoy et al., 2015; Sullivan et al., 2015). To test whether the measured increase in aspartate upon MPC inhibition could be responsible for the activation of quiescent NSPCs, we treated NSPCs with 5mM aspartate during quiescence induction and observed a significant increase in the number of proliferating, pH3 positive cells. Moreover, simultaneous treatment with UK5099 and aspartate further increased the ratio of pH3 positive NSPC cells (Fig 2F and Suppl Fig 2H).

Taken together, these data show that inhibition of MPC causes an increase in aspartate levels in quiescent NSPCs, which could result from increased GOT activity and/or increased import. Importantly, this increase in aspartate is sufficient to drive NSPC activation.

### MPC-deficient NSPCs can generate neurons through a shift in their metabolism

Once activated, NSPCs can proliferate and eventually differentiate into neurons. We therefore next tested the effect of UK5099 in already active/proliferative NSPCs. Similar to quiescent NSPCs, active NSPCs showed decreased OCR and increased ECAR upon treatment with UK5099 (Fig 3A), suggesting that these cells also use pyruvate to drive OXPHOS, despite lower levels of MPC1 and MPC2 than quiescent NSPCs (Fig 1E). Along with these findings, we also observed a significant decrease of TCA intermediates (Fig 3C) and a shift towards increased production of aspartate and asparagine with UK5099 treatment (Fig 3C), while no changes in the intermediates of glycolysis were found (Fig Suppl 3A). However, despite these changes in mitochondrial metabolism, UK5099 did not alter the proliferation rate of already active NSPCs (Suppl Fig 3B).

**Figure 3:**
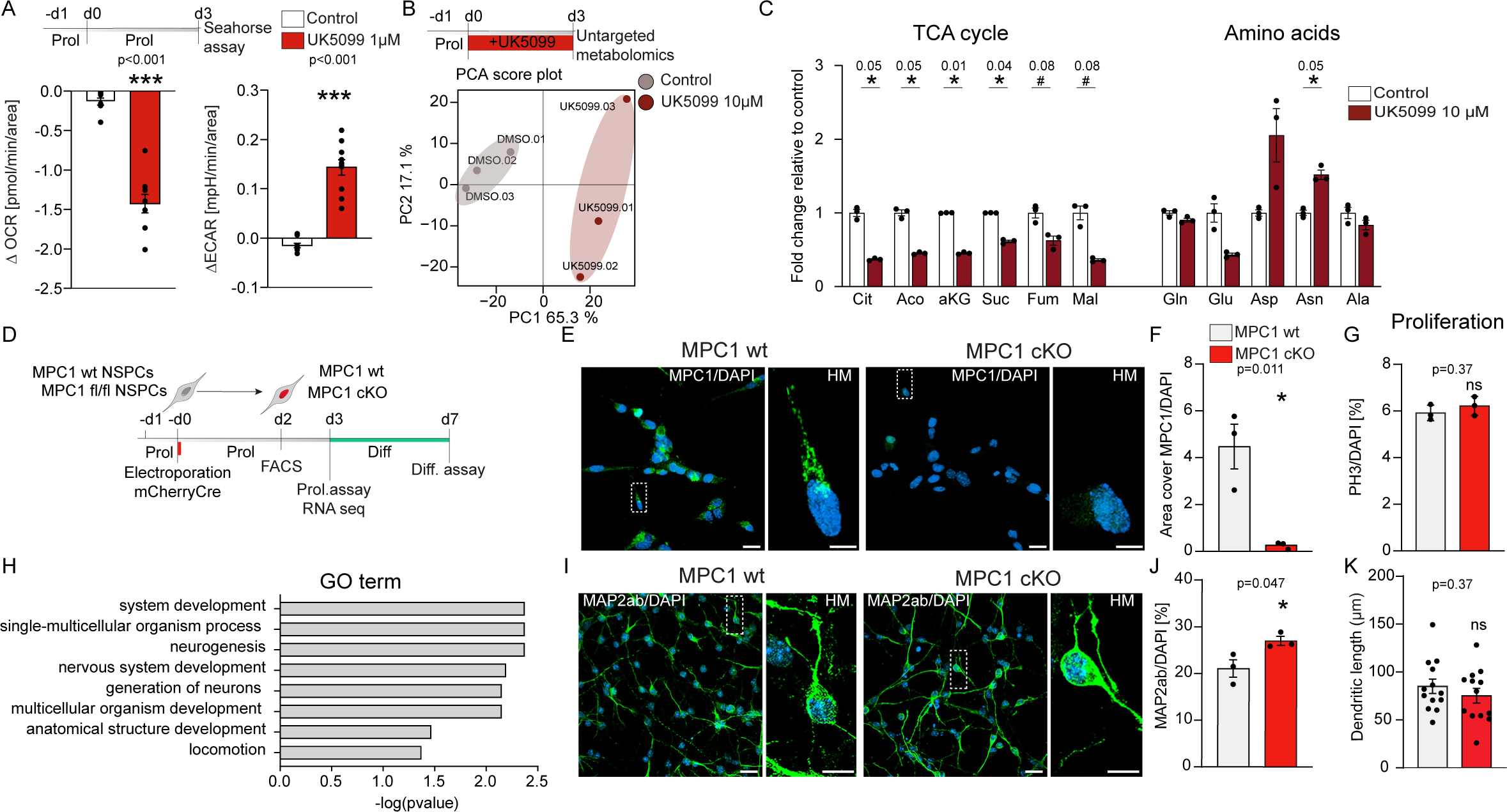
NSPCs lacking mitochondrial pyruvate important can still generate neurons through a shift in their metabolism. A. Experimental outline of seahorse analysis in proliferative NSPCs. Left graph represents OCR measurements in control and 1 μM UK5099 treated proliferative NSPCs. Right graph represents ECAR measurements in control and 1 μM UK5099 treated proliferative NSPCs. Bars represent mean ± SEM. ***p<0.001, n=9 wells from 2 independent experiments. (unpaired Student’s t-test). B. Top: Experimental outlines of untargeted metabolomics in proliferative NSPCs. Bottom: Principal analysis component of untargeted metabolomics from control and UK5099-treated proliferative NSPCs. Grey represents control group and red represents UK5099 treated cells. N= 4 biological replicates. C. Relative intensity of selected metabolites in control and UK5099 treated cells. Bars represent mean ± SEM. * adjusted p-value <0.05, # adjusted p-value <0.1, N=3 biological replicates. Cit: Citrate, Aco:Aconitate, aKG: alpha-ketoglutarate, Suc: Succinate, Fum: Fumarate, Mal: Malate, Gln: Glutamine, Glu: Glutamate, Asp: Aspartate, Asn: Asparagine, Ala: Alanine. D. Illustration shows the MPC1 knock-out system in cultured NSPCs. MPC1 wt/wt and MPC1 fl/fl NSPCs were electroporated with pCAGmCherryCre plasmid and selected by mcherry signal using fluorescent activated cell sorting (FACS) E. Representative low magnification (LM) and high magnification (HM) confocal images of MPC1 (green) and DAPI (blue) immunostaining in MPC1 wt and MPC1 cKO NSPCs after cre induction. Scale bar LM: 20 μm, HM: 5 μm F. Quantification of MPC1 intensity in MPC1 wt (grey) and MPC1 cKO (red) NSPCs. *p<0.05 n=3 electroporations, data show mean ± SEM, unpaired Student’s t-test. G. Quantification of pH3+ in MPC1 wt (gray) and MPC1 cKO (red). Data represent mean ± SEM. Not significant (ns), n=3 electroporations (unpaired Student’s t-test). H. GO term analysis of the upregulated genes in MPC1-cKO NSPCs. I. Representative low magnification (LM) and high magnification (HM) confocal images of neurons (MAP2ab, green) and DAPI (blue) in MPC1 wt and MPC1 cKO differentiated cells. Scale bar LM: 20 μm, HM: 5 μm J. Quantification of MAP2ab positive cells in MPC1 wt and MPC1 cKO cells. *p<0.05, n=3 electroporations, mean ± SEM, unpaired Student’s t-test. K. Quantification of the dendritic length of MAP2ab positive cells in MPC1 wt and MPC1 cKO cells *in vitro*. Not significant (ns), n=13 neurons per group (unpaired Student’s t-test).

Neuronal differentiation is associated with an increased expression of genes involved in OXPHOS (Beckervordersandforth et al., 2017; Shin et al., 2015). As glucose- or lactate-derived pyruvate is thought to be the major fuel for OXPHOS during differentiation, MPC-mediated pyruvate import should thus also play an important role during NSPC differentiation. We therefore tested if active NSPCs could differentiate into neurons in the presence of UK5099. We found a clear reduction in the total number of cells after 7 days of differentiation in the presence of UK5099 (Suppl. Fig. 3C), indicating that mitochondrial pyruvate might regulate cell survival or cell cycle exit during the differentiation process. However, surprisingly, UK5099-treated NSPCs were still able to give rise to microtubule associated protein 2ab (MAP2AB) positive neurons in a similar ratio to total cell numbers as untreated NSPCs (Suppl Fig 3D and E). These data highlight a large metabolic flexibility of NSPCs, which seem to use other metabolic pathways to sustain their energy needs for proliferation and differentiation when pyruvate import into mitochondria is disrupted.

To confirm these important findings, we generated an *in vitro* system of MPC1 knockout NSPCs. We isolated and expanded NSPCs from MPC1 fl/fl mice and wildtype control mice *in vitro* under proliferating conditions and induced Cre-mediated recombination through electroporation of a Cre-expressing plasmid. Three days after Cre-mediated deletion of MPC1 (Fig. 3D, Suppl. Fig. 3F), MPC1 protein levels were strongly reduced in proliferating MPC1 cKO NSPCs compared to MPC1 wt NSPCs (3E and F). Similar to UK5099-treated active NSPCs, MPC1 deletion did not affect proliferation of activate NSPCs (Fig 3G). While we also found a similar reduction in total cell numbers as in UK5099 treated NSPCs after 7 days of differentiation (Suppl. Fig. 3G), the deletion of MPC1 in active NSPCs did not affect the differentiation and the maturation into MAP2AB positive neurons. Interestingly, MPC1-KO NSPCs generated even slightly more MAP2AB positive neurons per total number of cells compared to control NSPCs (Fig. 3I, J and K).

To get further insights into the increased neuron production, we next performed RNA-Seq on active MPC1 wt and MPC1 cKO NSPCs. Commonly used NSPC markers were comparable between the two groups (Suppl. Fig. 3H, I and J), indicating that MPC1 cKO cells retained the main features of NSPCs despite the lack of MPC1. Among the differentially expressed genes, 160 genes were upregulated and 34, including Mpc1, were downregulated in the MPC1 cKO NSPCs (Suppl. Fig. 3K). Interestingly, gene ontology (GO) analysis of the upregulated genes showed a clear enrichment of GO terms involved in neuronal differentiation (Fig. 3H, Suppl. table 1), while the downregulated genes did not enrich in a specific GO category. Altogether, these results indicate that blocking the import of pyruvate into the mitochondria for OXPHOS does not affect neuronal differentiation of NSPCs, contrary to what would have been predicted.

### MPC1 deletion increases neurogenesis in vivo

We next tested whether deletion of MPC1 in adult NSPCs *in vivo* would affect their differentiation capacity 30 days after the first tamoxifen injection (Fig 4B). Cre-mediated MPC1 deletion at P30 led to a significant reduction in MPC1 mRNA and protein in the hippocampus, as shown by quantitative RT-PCR and western blot on whole hippocampal tissue of MPC1 cKO-tdTom and control mice at P60 (Suppl. Fig. 4A ,B and C). MPC2 was also downregulated (Suppl. Fig. 4C), as has been previously shown in other MPC1 knockout cell types (Bender et al., 2015; Schell et al., 2014, 2017; Vacanti et al., 2014; Vigueira et al., 2014)

**Figure 4:**
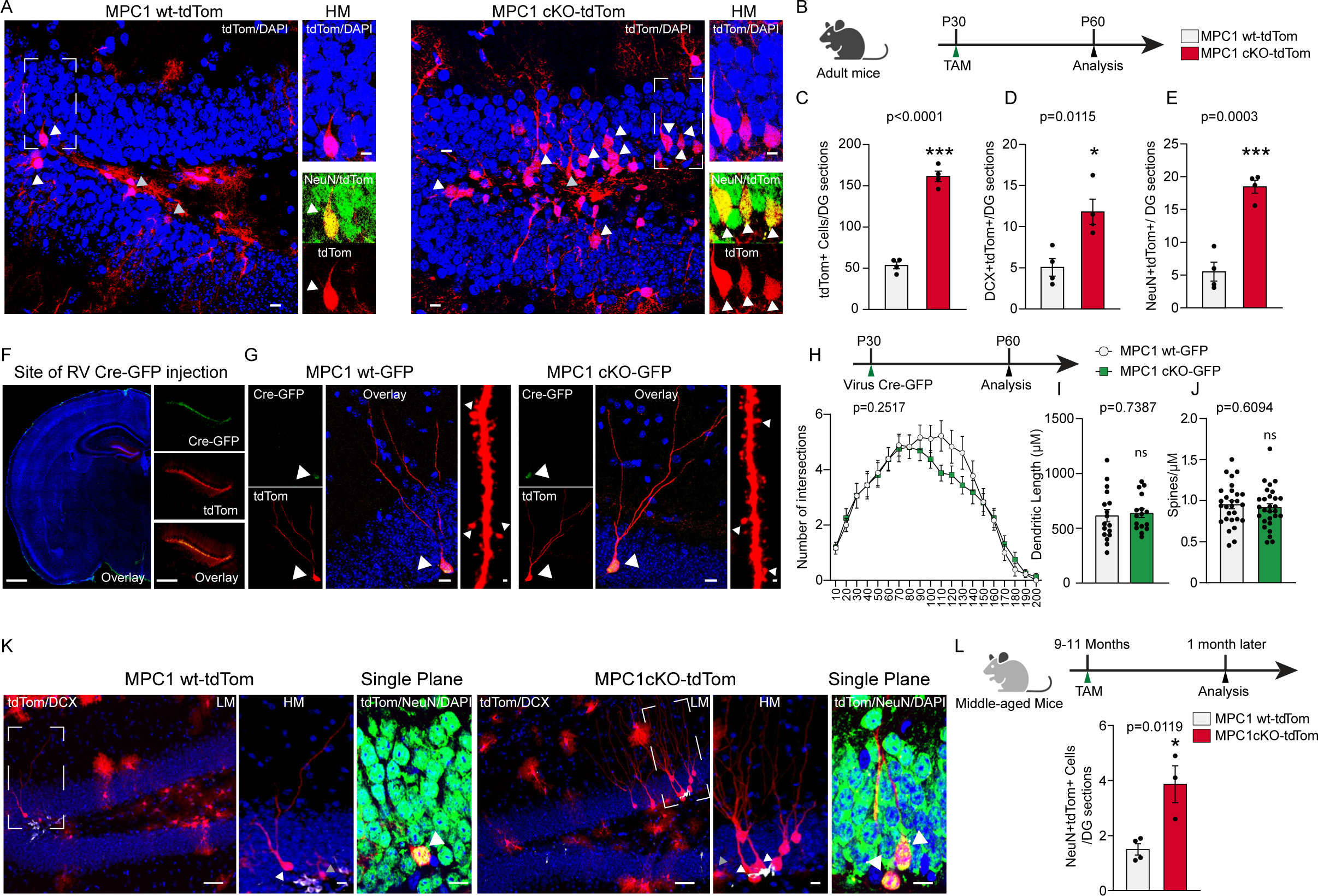
MPC1 deletion in NSPCs *in vivo* leads to increased numbers of progeny and increases neurogenesis. A. Representative low magnification (LM) and high magnification (HM) confocal images of TAM-induced tdTom recombination (tdTom, red), NeuN (green) and DAPI (blue) immunostaining at P60 in the DG of control MPC1 wt-tdTom and MPC1 cKO-tdTom mice. Scale bars: 10 μm. B. Experimental scheme. TAM injection at P30 (green arrow) and data analysis at P60. Below: Quantification of the recombined tdTom+ cells in the DG of control MPC1 wt-tdTom (grey) and MPC1 cKO-tdTom (red) mice at P60. C. The graph shows the average number of tdTom+ cells per DG sections (n=12 sections per mouse). Data are shown as mean ± SEM. ***p<0.001, n=4 mice per group (unpaired Student’s t-test). D. Quantification of the recombined DCX+dTom+ cells in the DG of control MPC1 wt-tdTom (grey) and MPC1 cKO-tdTom (red) mice at P60. The graph shows the average number of DCX+tdTom+ cells per DG sections (n=4 sections per mouse). Data show mean ± SEM. *p<0.05, n=4 mice per group (unpaired Student’s t-test). E. Quantification of the recombined NeuN+dTom+ cells in the DG of control MPC1 wt-tdTom (grey) and MPC1 cKO-tdTom (red) mice at P60. The graph shows the average number of NeuN+tdTom+ cells per DG sections (n=10 sections per mouse). Data show mean ± SEM. ***p<0.001, n=4 mice per group (unpaired Student’s t-test). F. Representative images of in vivo injections of retrovirus (RV) Cre-GFP in the DG. Scale bars: 250 μm. G. Representative confocal images of Cre-induced td-Tomato positive newborn neurons in the DG of MPC1 wt-GFP and MPC1 cKO-GFP mice. Arrowheads indicate the colocalization between nuclear GFP and tdTom signal. Scale bars: 10 μm. Representative high magnification (HM) confocal image of spines of newborn neurons in MPC1 wt-GFP and MPC1 cKO-GFP mice. Scale bars: 1 μm. H. Top: experimental scheme. Retro Virus Cre-GFP (RV Cre-GFP) injection at P30 (green arrow) and data analysis at P60 (black arrow). Below: Sholl analysis of dendritic complexity of newborn neurons in MPC1 wt-GFP and MPC1 cKO-GFP mice. n=4 mice per group, (unpaired Student’s t-test of AUC). I. Quantification of the dendritic length of newborn neurons in MPC1 wt-GFP and MPC1 cKO-GFP mice. Not significant (ns), n=4 mice per group (unpaired Student’s t-test). J. Spine density of newborn neurons in MPC1 wt-GFP and MPC1 cKO-GFP mice. Not significant (ns), n=4 mice per group, (unpaired Student’s t-test). K. Representative LM,HM and a single plan confocal images of TAM-induced td-Tomato recombination (red), DCX (white), NeuN (green) and DAPI (blue) immunostaining in the DG of middle-aged MPC1 wt-tdTom (grey) and MPC1 cKO-tdTom (red) mice. Scale bars: 10 μm. L. Top: Experimental scheme. TAM injection at 9-11 months (“middle-aged mice”) (green arrow), and data analysis after one month (black arrow). Below: Quantification of the recombined NeuN+dTom+ cells in the DG of control MPC1 wt-tdTom (grey) and MPC1 cKO-tdTom (red) mice. The graph shows the average number of NeuN+tdTom+ cells per DG sections (n=10 sections per mouse). Data show mean ± SEM. *p<0.05, n=3-4 mice per group (unpaired Student’s t-test).

Immunohistological analyses showed that the overall numbers of tdTom positive (tdTom+) cells in the DG were significantly increased in MPC1 cKO-tdTom mice compared to control mice one month after MPC1 deletion (Fig. 4A and C). This increase was due to increased numbers of neuroblasts and immature neurons marked by DCX+/tdTom+ (Fig. 4D, Suppl Fig 4D), as well as increased numbers of mature newborn neurons (NeuN+/tdTom+) (Fig. 4A and E). These data show that the generation of newborn neurons *in vivo* is not impaired by MPC1 deletion in NSPCs. On the contrary, the lack of MPC1 in NSPCs even led to a significant increase in the generation of newborn neurons.

To address whether the increased neurogenesis was due to a cell-autonomous MPC1 knockout effect in already activated NSPCs, we performed stereotaxic injection of a retrovirus encoding a Cre-recombinase fused to a GFP fluorophore into the DG of mice carrying floxed or wildtype MPC1 alleles, crossed to the tdTom lineage tracing line (Fig. 4F). As previously reported, this viral Cre-delivery strategy targets only activated NSPCs (Tashiro et al., 2007). This approach results in MPC1-deficient NSPCs (MPC1 cKO-GFP) or control NSPCs (MPC1 wt-GFP) in an otherwise unaltered microenvironment, and all progeny are labeled with tdTom and GFP. One month after virus injection, mature neurons were found in both MPC1 cKO-GFP and MPC1 wt-GFP mice (Fig. 4G). Despite the lack of MPC1, the neurons in the MPC1 cKO-GFP mice had similar morphology to the neurons in MPC1 wt-GFP mice, as assessed by Scholl analysis and dendritic length measurements (Fig. 4H and I). Similarly, total spine numbers were not changed (Fig. 4J) and the maturity of the spines was normal (Suppl. Fig. 4E). These astonishing findings show that even with disrupted oxidation of glucose-derived pyruvate, NSPCs can give rise to normal mature newborn neurons. This suggests that NSPCs and their progeny display metabolic flexibility and can compensate for impaired pyruvate import.

### MPC1 deletion also increases neurogenesis with advanced age

Hippocampal neurogenesis drastically drops with age (Kempermann, 2015), with more than 4-fold reduction of cycling NSPCs at 9 months compared to 2 months of age (Abdallah et al., 2010). Several recent studies suggest that this is due to increased dormancy of NSPCs (Ibrayeva et al., 2021; Lugert and Taylor, 2011; Urbán et al., 2019). Interestingly, when querying scRNA-Seq data from a recent publication which compared dormant, resting and proliferating NSPCs *in vivo* (Harris et al., 2021), we found that MPC1 and MPC2 were highest in dormant cells, and lowest in proliferating NSPCs (Suppl. Fig. 4F). As MPC1 deletion leads to an activation of quiescent NSPCs in young adult mice (Fig. 1K-L), we next tested whether deletion of MPC1 would have similar effects on dormant NSPCs from older animals. MPC1 deletion was induced in 9-11 months old MPC1 cKO-tdTom mice and the number of newborn neurons was assessed 30 days later. The number of tdTom+ neurons was significantly increased in the aged MPC1 cKO tdTom mice compared to the MPC1 wt tdTom mice (Fig. 4K and L), suggesting that in mice with advanced aged, NSPCs are also activated upon MPC1 ablation, leading to an increase in neurogenesis.

## Discussion

One of the most important regulatory steps of adult neurogenesis is the decision of NSPCs to maintain quiescence or enter an active state. This decision results from intrinsic and extrinsic instructions, which are still not fully understood (Urbán and Cheung, 2021). We here show that the maintenance of quiescence in adult NSPCs is dependent on mitochondrial pyruvate metabolism and requires MPC activity. MPC, which imports pyruvate into the mitochondria, is highly expressed in adult NSPCs and its loss of function leads to an activation of quiescent NSPCs and increased neurogenesis. Interestingly, recent studies have shown increased proliferation upon MPC deletion in other stem cells such as intestinal- and hair follicle stem cells (Flores et al., 2017; Schell et al., 2017). Furthermore, tumor cells slowed down proliferation after overexpression of MPC (Bensard et al., 2020; Schell et al., 2014), suggesting that there might be a common regulation of cellular activity by MPC.

Given the general idea that stem cells are mostly glycolytic, whereas their differentiated progeny use OXPHOS, the involvement of MPC in the quiescence of NSPCs shown here is surprising, as this carrier is generally considered a marker of oxidative metabolism. Our results suggest that quiescent type-1 cells display a more oxidative metabolism than commonly thought and the import of pyruvate into the mitochondria is necessary to maintain their quiescent state. Accordingly, we and other have reported that these cells have an elaborate mitochondrial network and can use fatty acids to fuel OXPHOS (Beckervordersandforth et al., 2017; Iwata et al., 2020; Khacho et al., 2016, 2019; Knobloch et al., 2017; Wani et al., 2022). A possible explanation for these discrepancies comes from a recent study comparing proteomic and transcriptomic levels of several metabolic proteins: Wani and colleagues could demonstrate that changes in TCA, FAO and OXPHOS proteins were poorly reflected at the transcriptional levels, while glycolytic enzymes showed better correlation between protein and mRNA expression (Wani et al., 2022).

The detailed molecular mechanisms by which mitochondrial pyruvate controls type-1 stem cell proliferation remain to be elucidated. The inhibition of pyruvate transport into the mitochondria has been shown to induce a metabolic shift towards oxidation of other substrates, in particular amino acids, which in turn rewire the metabolism of the cells (Divakaruni et al., 2017; Vacanti et al., 2014; Yang et al., 2014). Indeed, we also observe major metabolic changes when blocking mitochondrial pyruvate import, which lead to the activation of quiescent NSPCs. We show that the inhibition of MPC induces an increase of intracellular aspartate, despite a significant decrease in OCR and TCA metabolites. Aspartate is a key amino acid for cell proliferation, not only for protein synthesis, but also for purine nucleotide and pyrimidine nucleotide synthesis (Sullivan et al., 2015). Exogenous aspartate is sufficient to sustain proliferation even in the absence of a functioning ETC (Sullivan et al., 2015) and is also a limiting metabolite for cancer cell proliferation (Garcia-Bermudez et al., 2018). Furthermore, aspartate levels have been shown to directly influence hematopoietic stem cell proliferation (Qi et al., 2021). Our findings that the addition of exogenous aspartate mimics the effect of UK5099 on quiescence exit of NSPCs are thus in line with these previous findings. The additive effect of aspartate and UK5099 suggest that both mechanisms, aspartate uptake and/or aspartate biosynthesis, contribute to the intracellular pool of aspartate. Despite its clear effect on NSPC activation, it remains to be clarified how exactly increased aspartate is exerting its effect. A possible explanation is that aspartate is used for the synthesis of purine/pyrimidine and asparagine, similar to findings in hematopoietic stem cells (Qi et al., 2021). Indeed, our metabolomic and mRNA expression data point in this direction, showing an increase of both asparagine synthetase and asparagine synthesis in quiescent NSPCs upon MPC inhibition.

A shift in the balance of NSPC quiescence and activation is expected to affect net neurogenesis, as the production of new neurons is the default fate of activated adult hippocampal NSPCs (Denoth-Lippuner and Jessberger, 2021). Accordingly, we found a substantial increase in the number of newborn neurons upon MPC deletion. However, given the commonly accepted view that most neurons heavily rely on glucose- and lactate-derived pyruvate for OXPHOS (Ashrafi and Ryan, 2017; Magistretti and Allaman, 2018) such an increase in morphologically normal neurons is astonishing, as one would expect that newborn neurons cannot mature properly without pyruvate oxidation. These results suggest that newborn neurons might have a higher metabolic flexibility than previously thought if their preferential metabolic pathway is blocked. In line with this, it has been shown that several types of mature neurons are indeed able to sustain their functions by shifting their metabolism to other substrates, which can fuel OXPHOS when MPC is inhibited (DeLaRossa et al., 2022; Divakaruni et al., 2017; Ghosh et al., 2016). Indeed, we show that MPC1 ckO NSPCs are able to generate mature neurons *in vitro* and *in vivo*. These findings highlight a new form of metabolic plasticity used by NSPCs to differentiate into mature neurons. Interestingly, MPC1 knockout seems to prime NSPCs towards neuronal differentiation, as our RNA-Seq data suggest. These data illustrate that the metabolic state might directly influence fate decision. Whether this neuronal priming is achieved via epigenetic modifications in MPC1 knockout NSPCs remains to be determined.

Hippocampal neurogenesis decreases strongly with age, due to terminal differentiation and increased deep quiescence of NSPCs (Lugert and Taylor, 2011; Urbán et al., 2019). Remarkably, our data show that MPC deletion is also sufficient to wake-up these more dormant NSPCs in aged mice, resulting in increased number of newborn neurons. Supporting these findings, we found that MPC is highest in dormant NSPCs in a recent scRNA-Seq study comparing different states (Harris et al., 2021), which is also in line with our *in vitro* quiescence data.

In conclusion, we have shown that quiescence of NSPCs is an actively maintained state, requiring MPC-mediated import of pyruvate into the mitochondria. MPC inactivation leads to activation of quiescent NSPCs and subsequently increased neurogenesis even in aged mice. Our findings thus describe a new mechanism through which mitochondrial metabolism controls NSPC function.

## Material and Methods

### Animals

All studies were approved by the services de la consummation et des affaires veterinaires (SCAV) of Geneva in Switzerland. Mice were group-housed with littermates in standard housing on a 12:12 h light/dark cycle. hGFAP-CreERT2 mice (Hirrlinger et al., 2006) have been obtained from Professor Nicolas Toni (Center for Psychiatric Neurosciences, Lausanne University Hospital, CHUV), MPC1fl/fl mice (Gray et al., 2015) from Professor Eric Taylor (University of Iowa) and tdTom fl-STOP-fl (Ai14, (Madisen et al., 2010)) from Professor Ivan Rodriguey (University of Geneva). All mice were on a C57BL/6 background.

Genotyping was performed on DNA extracted from phalange biopsies using the following primers: hGFAP-CreERT2: F-CAG GTT GGA GAG GAG ACG CAT CA, R-CGT TGC ATC GAC CGG TAA TGC AGG C; MPC1fl/fl: F1-CCT ATT CTC TAG AAA GTA TAG GAA CTT CGT CGA, F2-GTG AGC CCA GAG CTA CGA AGG ATC GGC, F3-GGA AAG AAA AAG GTG TCC AAT TTT AGC TCT GCA; tdTom fl STOP-fl: F 5’-CTG TTC CTG TAC GGC ATG G-3’,R 5’-GGC ATT AAA GCA GCG TAT CC-3’, tdTom wt/wt:-F 5’-AAG GGA GCT GCA GTG GAG TA-3’,R 5’-CCG AAA ATC TGT GGG AAG TC-3’.

### *In vivo* treatments

Tamoxifen (Sigma, 100 mg/kg) was injected intraperitoneally (i.p) for 5 consecutive days (P30 to P35) to induce recombination *in vivo.* The tamoxifen was dissolved in sunflower oil (S5007 Sigma-Aldrich)

### Analysis of transcriptomic resources

We queried available single cell RNA-Seq data for MPC expression (Hochgerner et al., 2018; Shin et al., 2015) For the expression of MPC1 in NSPCs, the original unsupervised clustering and pseudotime subgroups (S1 to S5) from Shin et al. were further divided in 3 sub-groups: S1-S2, S3 and S4-S5 reflecting respectively a more quiescent to proliferative state.

### Cell culture

Adult mouse neural stem/progenitor cells (NSPCs) from hippocampus of 7 week-old C57/Bl6 (MPC1 wt) or MPC1 fl/fl (MPC1 cKO) male mice were isolated as previously described (Ramosaj Madsen et al). For propagation, cells were cultured as spheres in DMEM/F12 GlutaMAX (#31331-028, Invitrogen) supplemented with N2 (#17502048, Invitrogen), human EGF (20 ng/ml, #AF-100-15, Peprotech), human FGF-basic (20 ng/ml, #100-18B, Peprotech), heparin (5 μg/ml, #H3149-50KU, Sigma). Medium contained as well an antibiotic-antimycotic (#15240062, Invitrogen). Medium was changed every second day.

For proliferation experiments, cells were plated (42’000 cells/cm^2^ ) on glass coverslip (#10337423, Fisher Scientific) coated with Poly-L-ornithine (50 μg/ml, #P36655, Sigma) and laminin (5 μg/ml, #L2020, Sigma) and grown as described above. Quiescence was induced as previously described(Knobloch et al., 2017). Proliferating NSPCs (40’000 cells / cm^2^) were plated on coated glass coverslip or coated plastic cell culture well-plate. Twenty-four hour later, medium was change to quiescence medium based on DMEM/F12 GlutaMAX, supplemented with N2, 20 ng/ml of human FGF-basic, 5 μg/ml heparin and 50 ng/ml BMP-4 (#5020BP, R&D System). Seventy-two hours after induction, cells were fully quiescent and used for subsequent experiments. For UK5099 experiment, quiescence was fully established for 3 days, then cells were washed off, trypsinized and plated (220’000 cells/cm^2^) on coated coverslips in fresh quiescence medium

For comparison, proliferative NSPCs were plated at a lower density (13’000 cells/ cm^2^) under proliferative conditions. The quiescence state was reversed by collecting quiescent NSPCs and replating them in proliferative conditions (50’000 cells/cm^2^) for 3 days (Fig 2A).

For differentiation experiments, cells were plated on glass coverslip (#10337423, Fisher Scientific) or cell culture plates coated with Poly-L-ornithine (50 μg/ml, #P36655, Sigma) and laminin (5 μg/ml, #L2020, Sigma) and cultured with a medium containing 1/5 of human EGF and human FGF-basic, i.e 2 ng/ml. After 3 days, medium was change to medium without any human EGF and human FGF-basic. Cells were fixed for 15-20min with 4% paraformaldehyde (PFA) after 6-7 days of differentiation. Cells were washed 2x for 10min with PBS and stored at 4°C prior staining.

### Treatment with UK5099 and aspartate

Proliferative NSPCs were treated with either 1 μM or 10 μM of UK5099 (#PZ0160, Sigma) for 72 hours. For quiescence experiments NSPCs were either treated with 1 μM or 10 μM of UK5099 (#PZ0160, Sigma) simultaneously with quiescence induction or after establishment of quiescence. For differentiation experiments, proliferative NSPCs were treated with 1 μM or 10 μM of UK5099 for 48h. Then cells were washed off and plated (65000 cells/cm^2^) on coated coverslips in media containing 1/5 of the growth factors and indicated UK5099 concentration.

Similarly to UK5099 treatment, 1 mM, 2.5 mM or 5 mM Aspartate (#1690.2, Roth) was used during the induction of quiescence.

### Metabolic Fuel Flux Assays

Oxygen consumption rate (OCR) and extracellular acidification rate (ECAR) were measured with the Seahorse XFe96 Analyzer (Agilent). 4000 NSPCs/well were plated on 96-well seahorse XF96 cell culture microplates with proliferative medium. After 24h, quiescence was induced (see cell culture section). After three full days of quiescence condition, cells were carefully washed with unbuffered DMEM (#103575-100, Agilent) supplemented with 10 mM glucose (#103577-100, Agilent), 2mM glutamine (#103579-100, Agilent) and 1 mM pyruvate (#103578-100, Agilent), and a final volume of 180 μl assay media was added to cells. The plate was allowed to equilibrate in a non-CO_2_ incubator at 37°C for one hour prior to analysis

Mito Fuel Flex Test (#10360-100, Agilent) was performed according to manufacturer’s protocol. Baseline OCR was monitored for 14min followed by different combinations of inhibitor injections as reported in Table 1 with OCR measurements. The Mito Fuel Flex test inhibits the import of the three main metabolic substrates: pyruvate, glutamine and fatty acids using UK5099 (2 μM, #PZ0160, Sigma), BPTES (3 μM, #SML0601, Sigma) and etomoxir (4 μM, #E1905, Sigma), respectively.

**Table 1:** Combination of inhibitors.

| Metabolic test | Injection n°1 | Injection n°2 |
| --- | --- | --- |
| Pyruvate dependency | UK5099 | Etomoxir + BPTES |
| Pyruvate capacity | Etomoxir + BPTES | UK5099 |
| Glutamine dependency | BPTES | UK5099 + Etomoxir |
| Glutamine capacity | UK5099 + Etomoxir | BPTES |
| Fatty acid dependency | Etomoxir | UK5099 + BPTES |
| Fatty acid capacity | UK5099 + BPTES | Etomoxir |

Dependency was calculated as:

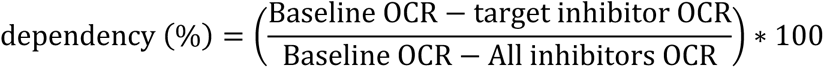

Capacity was calculated as:

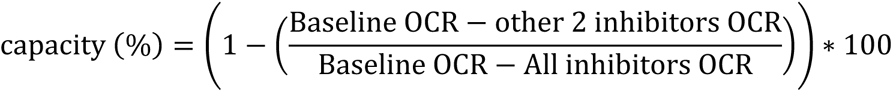

Flexibility was calculated as:

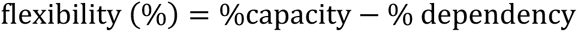

To assess the effect of MPC1 inhibition, baseline OCR and ECAR were measured after 14min in quiescent NSPCs. Different combinations of drugs: UK5099 (1 μM), etomoxir (20 μM), BPTES 3 (μM), Etomoxir/ BPTES, UK5099/Etomoxir or UK5099/BPTES were injected and OCR and ECAR were measured over 40min. The change in OCR or ECAR were calculated as (ΔOCR) = (OCR following drug injection−OCR at baseline) and (ΔECAR) = (ECAR following drug injection−ECAR at baseline).

In order to normalize each well to cell numbers, after the run, cells were incubated with Hoechst (1:2000, #H1399, Sigma) for 15min and fixed with 4% PFA for 15min. Cells were then washed with PBS and all the wells were imaged with a Thunder microscope (Leica, DMi8). Hoechst signal was thresholded and the area cover was measured using the software Fiji (Image-J, 2.0.0).

### Eletroporation and FACS separation

Proliferating MPC1 wt and MPC1 cKO NSPCs were tranfected with a plasmid encoding mCherry fused to Cre-recombinase (7 μg/μl, pCAGmCherryCre), using a 4D-Nucleofector X Unit EA (#LZ-AAF-1002X, RUWAG) and the P3 primary cell 4D-Nucleofector X solution (#LZ-V4XP-3024, RUWAG). NSPCs were cultured in normal proliferative medium for 48h before being subjected to Fluorescence Activated Cell Sorting (FACS). Cells were trypsinized and resuspended in EDTA-DPBS (#E8008, Merck Millipore) and kept in ice until the sorting. mCherry positive cells were sorted on a MoFlo Astrios EQ cell sorter (Beckman Coulter) and collected in proliferative medium or directly frozen on dry ice.

### Immunocytochemistry

To visualize MPC1, cells plated on glass coverslip were fixed with 4% PFA and 0.2% Glutaraldehyde for 15min. For all the other stainings, cells were fixed with 4% PFA for 15-20min. Cells were washed 2x for 10min with PBS and stored at 4°C prior staining. Cells were blocked for 45min with blocking buffer containing 0.2% Triton-X (#X100, Sigma), 3% Donkey serum (#S30, Sigma) in Tris-buffered saline (TBS, 50 mM Tris-Cl, pH 7.4, 150 mM NaCl) and incubated with the indicated primary antibodies diluted in blocking buffer at 4°C overnight: mouse-pH3 (1:2000, Abcam, ab14955), rabbit-Ki67 (1:500, Abcam, ab15580), rabbit-MPC1 (1:100, Sigma-Aldrich, HPA045119), mouse-Map2b (1:500, Sigma-Aldrich, M2320). Cells were washed 3x in TBS for 10min and incubated with secondary antibodies in blocking buffer for at least 1h at room temperature, protected from light (AlexaFluor; donkey anti-mouse 488, donkey anti-rabbit 488, donkey anti-rabbit 647,1:250). Cells were washed 2x 10min with TBS. Nuclei were stained with DAPI (1:5000, #D9542, Sigma) for 5min and washed 2x with TBS. Coverslips were mounted with a self-made PVA-DABCO-based mounting medium.

For mitochondrial visualizations, living cells were incubated with 200nM Mitotracker Deep Red FM (#M22426, Invitrogen) for 15min at 37°C. Cells were fixed and immunolabelled as mentioned above.

### Image acquisition and analyses

All images used to assess the ablation of MPC1, the quantification of MAP2AB and the mitochondrial shape in MPC1 wt and MPC1 cKO were acquired with a confocal microscope (Zeisse, LSM780) with a 40x or 63x objectives. Images were analysed using the software Fiji (ImageJ 2.0.0). Dentritic length was measured using the plugin “Simple Neurite Tracker” on Fiji.

All images to assess the proliferation were acquired in a blinded manner with an epifluorescent microscope (Nikon 90i or Leica DMi8) with a 20x or 40x objective. Analyses of the images was always done in a blinded manner.

For DAPI area cover, 56 tiles with a 10X objectives were acquired with an epifluorescent microscope (Leica DMi8). Area over was measured in thresholded images.

For mitochondrial analysis, serial z-stacks (0.8 µm) were taken using a confocal microscope (Zeisse, LSM780) with a 63x objective.

Analysis of mitochondria was performed using the software Imaris (9.8.0, Bitplane) as previously described (Zehnder et al., 2021). Surfaces were created on the mitotracker Deep Red signal. Sphericity values were exported as Excel files. Mitochondrial length was measured using the freehand tool in Fiji as previously described (Iwata et al., 2020).

### RNA-Seq and qRT-PCR on cultured NSPCs

For RNA-Seq, MPC1 wt and MPC1 cKO NSPCs were tranfected with a plasmid pCAGmCherryCre and sorted according to mcherry (as described above). Directly after the sort cells were collected and frozen down prior RNA extraction. RNA was extracted using RNeasy micro kit (Qiagen) according to the manufacturer’s protocol. RNA-Seq data were generated and analyzed by Alithea Genomics SA (Switzerland). For *in vitro samples,* NSPCs were collected and snap frozen on dry ice. RNA was extracted using RNeasy mini kit (#74134, Qiagen) according to the manufacturer’s protocol, followed by cDNA synthesis using SuperScript IV system (#15327696, Invitrogen). RT-qPCR was performed using PowerSYBR Grenn PCR Master Mix (Fisher scientific, #10658255). The following primer were used:

*Mpc1*: Forward: 5’-GAC TAT GTC CGG AGC AAG GA-3’, Reverse: 5’-TAG CAA CAG AGG GCG AAA GT-3’. *Mpc2*: Forward: 5’-TGC TGC CAA AGA AAT TG-3’, Reverse: 5’-AGT GGA CTG AGC TGT GCT GA-3’. *Got1*: Forward: 5’-CTT TAA GGA GTG GAA AGG TAA C-3’, Reverse: 5’-GAG ATA GAT GCT TCT CGT TG-3’. *Got2*: Forward: 5’-TAT CAA AAA TCC CAG AGC AG-3’, Reverse: 5’-ATT CTT TTT CTT CAC CAC GG

-3’. *Asns*: Forward: 5’-CCA AGT TCA GTA TCC TCT CC -3’, Reverse: 5’-TAA TTT GCC ACC TTT CTA GC-3’. *Slc1a3*: Forward: 5’-TCA TCT CCA GTC TCG TCA CA-3’, Reverse: 5’-CAC CAC AGC AAT GAT GGT AGT A-3’. *β-actin*: Forward: 5’-GCC CTG AGG CTC TTT TCC AG-3’, Reverse: 5’-TGC CAC AGG ATT CCA TAC CC-3’. β-actin was used as a housekeeping gene to normalize the obtained CT values and each sample gene expression was calculated using the comparative ΔΔCT method.

### qRT-PCR on brain tissue

Total RNA from hippocampal tissue was isolated using trizol reagent (Thermo Fisher) method and RNA concentration was determined using NanoDrop One (Thermo Scientific). Reverse transcription was performed with 500-1000ng of DNase-treated (Promega, RQ1 M610A) total RNA using Go script RTase (Promega A6001). The qRT-PCR was done on a Bio-Rad CFX Connect^TM^ optics module. CT values were normalized to β-actin.

*Mpc1*: Forward: 5’-GAC TAT GTC CGG AGC AAG GA-3’, Reverse: 5’-TAG CAA CAG AGG GCG AAA GT-3’. *Mpc2*: Forward: 5’-TGC TGC CAA AGA AAT TG-3’, Reverse: 5’-AGT GGA CTG AGC TGT GCT GA-3’. Beta-Actin, Forward 5’ -GGCTGTATTCCCCTCCATCG-3’, Reverse 5’ CCAGTTGGTAACAATGCCATGT-3’

### Protein extraction and western blotting

Hippocampal brain and cellular extracts were homogenized in RIPA lysis buffer (50mM Tris HCl , pH 7.5, 150 mM NaCl, 1% NP-40, 0.5% sodium deoxycholate, 0.1 SDS, 1mM EDTA, 10% Glycerol, protease inhibitors), and resolved by SDS-PAGE in 8-15% polyacrylamide gels. The concentration of the proteins was determined by Bradford protein assay using bovine serum albumin as standard and analyzed by western blotting with specific antibodies.

Antibodies used: rabbit-MPC1 (Sigma, HPA045119), mouse-MPC2 (home-made), goat-VDAC (Santa Cruz, sc-8829), mouse-HSP70 (Invitrogen, MA3-028), total OXPHOS antibody cocktail (Abcam, MS604-300), anti-IgG-rabbit-HRP (Dako, P0217), anti-IgG-mouse-HRP (Dako, P0447) anti-IgG-Goat-HRP (Santa Cruz, sc-2304).

### Tissue preparation and immunohistochemistry

All experimental mice were deeply anesthetized with sodium pentobarbitone (6mg/100g body weight, i.p.) at different time points after tamoxifen injection (P33, P38, P60, 10-12 months) and immediately perfused intracardiacally with fresh 4% paraformaldehyde in 0.1M phosphate-buffered saline (pH 7.4). Brains were postfixed overnight at 4°C. Coronal sections of 50µm and 70µm were cut using a vibratome (Leica) and stored at 4°C in 1X PBS supplemented with 0.02% sodium azide. For immunohistochemical analysis, sections were permeabilized for 1h in PBS containing 0.3% Triton X-100, and 10% donkey or goat serum and then immunolabeled overnight at 4°C on an orbital shaker, using the following primary antibodies: mouse-NeuN (1:200, Merck-Millipore, MAB377), chicken-GFP (1:500, Abcam, ab13970), rabbit-DCX (1:400, Cell Signaling technology, 4604) guinea pig-DCX (1:100, Merk-Millipore, AB2253) and rabbit-Ki67 (1:400, Abcam, ab16667). After the primary antibody incubation, the sections were washed again three times in PBS for 5min and incubated for 2h at RT with fluorescent secondary antibodies (AlexaFluor, goat anti-mouse 647; donkey anti-mouse 647, goat anti-rabbit 488, donkey anti-rabbit 488 and donkey anti-guinea pig 647; donkey anti-goat 488, donkey anti-chicken 488, 1:300) diluted in PBS. Finally, nuclei were counterstained with 4’, 6-diamidino-2-phenylindole (DAPI) (Invitrogen, 1:10000) and then washed before mounting with a FluorSave^TM^ reagent (Merck-Millipore).

### Metabolomics Analysis

For untargeted metabolic profiling, adherent cells were washed with a solution containing 75 mM Ammonium Carbonate (#379999-10G, Sigma) and the pH was buffered at 7.4 using acetic acid. After removing the liquid, cells were frozen by holding the place on dry ice. Extraction of metabolites was performed by adding ice-cold acetonitrile (#34967, Sigma), methanol (#34966, Sigma) and water (#W6-500, Thermo Scientific) (4:4:2, v/v) solvent mixture to cells. The extracts were centrifuged at 13’000 rpm for 2 minutes prior analysis.

Non-targeted metabolomics analysis was performed by flow-injection – time-of-flight mass spectrometry on an Agilent 6550 QTOF system (PMID21830798). The instrument was set to scan in full MS at 1.4 Hz in negative ionization, 4 GHz High Res mode, from 50 to 1000 m/z. The solvent was 60:40 isopropanol:water supplemented with 1 mM NH4F at pH 9.0, as well as 10 nM hexakis (1H, 1H, 3H-tetrafluoropropoxy) phosphazine and 80 nM taurochloric acid for online mass calibration. The injection sequence was randomized. Data was acquired in profile mode, centroided and analyzed with Matlab (The Mathworks, Natick). Missing values were filled by recursion in the raw data. Upon identification of consensus centroids across all samples, ions were putatively annotated by accurate mass and isotopic patterns. Starting from the HMDB v4.0 database, we generated a list of expected ions including deprotonated, fluorinated, and all major adducts found under these conditions. All formulas matching the measured mass within a mass tolerance of 0.001 Da were enumerated. As this method does not employ chromatographic separation or in-depth MS2 characterization, it is not possible to distinguish between compounds with identical molecular formula. Principal analysis component and statistical comparison between the groups were performed using RStudio. Pathway enrichment analysis was performed using MetaboAnalyst 5.0

### Virus preparation

CAG-Cre-GFP virus was produced as previously described (Zhao et al., 2006). Briefly human embryonic kidney 293T cells were transfected with a pCAG-GFPcre, pCMV-gp and pCMV-vsvg using Lipofectamine 2000 (#10696153, Fisher Scientific) in Opti-MEM medium (#11520386, Fisher Scientific). Forty-eight hours post-transfection, virus was collected by filtering the cell culture medium through a 0.22 µm filter. The filtrate was then concentrated twice using ultra-centrifugation at 19400 rpm. The viral pellet was resuspended in PBS, and used for stereotaxic injections.

### Virus injections *in vivo*

One month old (P30) recombined MPC1cKO-tdTom and control MPC1 wt-tdTom mice were anesthetised using isoflurane at 5% (w/v), placed in a small animal stereotaxic frame (David Kopf Instruments) and maintained at 2.5% isoflurane (w/c) for the duration of surgery. Corneal and pinch reflexes were regularly tested to confirm anaesthetic depth. Lacryvisc (Aicon, Switzerland) was applied to prevent corneal drying and lidocaine applied topically to the skin overlying the skull. After exposing the skull under aseptic conditions, a small hole was drilled in the skull overlying the DG. Retro Virus Cre-GFP (RV Cre-GFP) was injected monolaterally into the DG from the bregma (AP -2 mm, L+1,5 mm, 2.3 DV mm from skull) at a rate of 100 nL/min-1 (1.5 µl total volume) using a Hamilton syringe and CMA400 pump (CMA System, Kista, Sweden). Mice were sacrified 4 weeks after virus injections.

### Confocal microscopy acquisition and image analysis

All images were collected on a Leica confocal imaging system (TCS SP8) with a 20x or a 63× oil immersion objective, Leica Thunder imaging system (DMi8) with 20X objective and Spinning Disk confocal imaging system (Nikon Ti2/ Yokogawa CSU-W1) with 60X and 100X oil immersion objectives. For the quantification of recombined tdTom positive cells with specific markers, serial sections starting from the beginning of the DG were used. Quantification of tdTom positive and Ki67 positive cells labelling with different markers DCX, NeuN, were performed using the software Fiji (ImageJ 2.0.0). To confirm Ki67, DCX and NeuN colocalization with tdTom, single optical sections at 63X magnification were used.

For Sholl analysis, the Cre-virus transfected GFP and tdTom double-positive newborn neurons were imaged with a 63X (0.75 NA) objective. Z-stacks were taken at 1µm intervals, and dendrites were traced using the Neurolucida software (version 10, mbs Bioscience). 15-18 neurons from 4 mice per group were analyzed. Dendritic spine density and spine morphology were assessed as previous described (Petrelli et al., 2020; Sultan et al., 2015) with little adjustments. Briefly, dendrites of 20-30 Cre-virus transfected GFP and tdTom double-positive newborn neurons were imaged using a 63x (2.5 NA) objective. The dendritic length and the number of spines were analysed using the software Fiji (ImageJ 2.0.0). Spine density was expressed as the number of the spines divided by dendritic length. Spine morphology was classified in two groups based on the maximal diameter of the spine head, as measured on maximal projections with Fiji (ImageJ 2.0.0). immature spines (thin spines) were defined as 0.25 to 0.6µm and mature spines (Mushroom) >0.6µm. The percentage of each type of dendritic spine was then expressed by neuron for each mice (20-30 neurons per mice, 4 mice per group). The data were expressed as the ratio between immature spines and mature spines. Images were analyzed in a blinded manner.

### Statistical Analyses

For two simple comparisons, unpaired t-test was used. For metabolomics data, p-value was adjusted for multiple comparisons. For comparisons with more than 2 groups, one-way ANOVA was performed followed by Holm-Sidak’s multiple comparison tests. For dendritic complexity in Sholl analysis, Area Under Curve (AUC) was calculated followed by unpaired t-test. All analyses were performed using GraphPad Prism 8.0.2 software.

### Illustration software

For illustration schemes, Biorender software (Biorender, 2021) and Adobe Illustrator (version 25.0, Adobe Inc) was used.

## Acknowledgements

We thank Eric Taylor (University of Iowa, USA) and Jarred Rutter (University of Utah Medical School, USA) who provided us with the MPC1 floxed mice, and Andrés de la Rossa, who initiated this project and generated the MPC1Flox;GFAP-Cre mice. We further thank Darcie Moore (University of Wisconsin, USA) and Sebastian Jessberger (University of Zurich, Switzerland) for helpful discussions on the manuscript. This work was supported by funding from the University of Geneva (SNSF #31003A_179421 to JCM) and from the University of Lausanne (to MK).

## Author contributions

F.P. and V.S conceived and performed experiments and analyzed and interpreted data. S.M. performed experiments. N.Z performed the metabolomic analysis. F.P, J.C.M and M.K. developed the concept, conceived experiments and interpreted data. F.P. ,V.S and M.K. wrote the manuscript, with input from all authors.

---

**Supplementary Fig 1:**
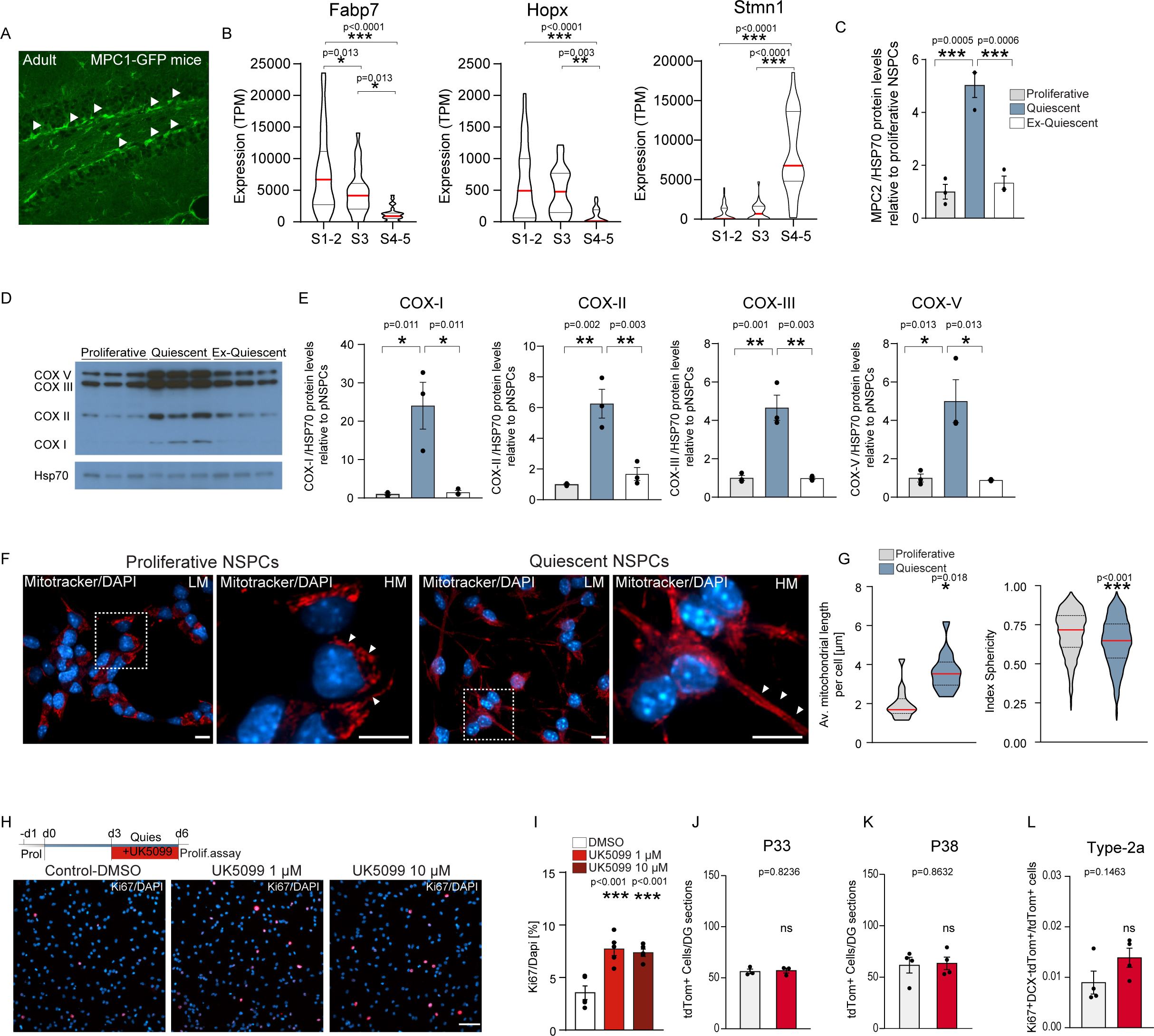
MPC is necessary to maintain the quiescence in NSPCs, related to figure 1. A. Representative sagittal confocal image (40X) of adult DG of MPC1-GFP mice (Gensat data, http://www.gensat.org/imagenavigator.jsp?imageID=26000) B. Violin plots of *Fabp7*, *Hopx* and *Stmn1* expression in quiescent (S1-2), more activated NSPCs (S3) and intermediate progenitor cells (S4-5). Data queried from Shin et al. Cell Stem Cell 2015. Red line represents mean. *p<0.05, **p<0.01, ***p< 0.001 (one-way ANOVA followed by post-hoc test) C. Quantification of MPC2 expression in proliferative (grey), quiescent (light blue) and ex-quiescent (grey) NSPCs. MPC2 expression was normalized to HSP70 levels and expressed as fold change to proliferative NSPCs. Bars represent mean ± SEM. ***p<0.001, n=3 biological replicates (one-way ANOVA followed by post-hoc test). D. Representative western blots of the different complexes of the oxidative phosphorylation chain: cytochrome c oxydase (COX)-I (18 kDa), COX-II (30 kDa), COX-III (48 kDa), COX-V (55 kDa) and HSP70 (70 kDa). E. Quantification of COX-I, COX-II, COX-III and COX-V in proliferative (grey), quiescent (light blue) and ex-quiescent (grey) NSPCs. Protein levels were normalized to HSP70 levels and expressed as fold change to proliferative NSPCs. Bars represent mean ± SEM.*p< 0.05 ,**p<0.01, n=3 biological replicates (one-way ANOVA followed by post-hoc test). F. Representative LM and HM confocal images of mitotracker (red) and DAPI (blue) immunostaining in proliferative and quiescent NSPCs. Arrowhead show mitochondria shape. Scale bars: 10 μm. G. Quantification of mitochondrial length and sphericity in proliferative and quiescence NSPCs. *p<0.05, ***p<0.001. n=13-14 cells from 3 CS. (Mann-Whitney test). H. Top: Experimental outline of UK5099 treatment after quiescence establishment. Bottom: Representative images of Ki67 (red) and DAPI (blue) immunostaining in quiescent NSPCs treated with UK5099. Scale bar: 50 μm. I. Quantification of Ki67+ in DMSO (white) or UK5099 (light and dark red) treated quiescent NSPCs. Data represent mean ± SEM. ***p<0.001. n=5 CS from 2 independent experiments (one-way ANOVA followed by post-hoc test) J. Quantification of the recombined tdTom positive (tdTom+) cells in the DG of MPC1 cKO-tdTom (red) and control MPC1 wt-tdTom (grey) mice at P33. The graph shows the average number of tdTom+ recombined cells per DG sections (n=10 sections per mouse). Data show mean ± SEM. Not significant (ns), n=3 mice per group (unpaired Student’s t-test). K. Quantification of the recombined tdTom+ cells in the DG of and control MPC1 wt-tdTom (grey) and MPC1 cKO-tdTom (red) mice at P38. The graph shows the average number of tdTom+ recombined cells per DG sections (n=10 sections per mouse). Data show mean ± SEM. Not significant (ns), n=4 mice per group (unpaired Student’s t-test). L. Quantification of type-2a Ki67+ DCX-tdTom+ cells on total number of tdTom+ cells in the DG of control MPC1 wt-tdTom (grey) and MPC1 cKO-tdTom (red) mice. Data represent mean ± SEM. Not significant (ns), n=4 mice per group (unpaired Student’s t-test).

**Supplementary Figure 2:**
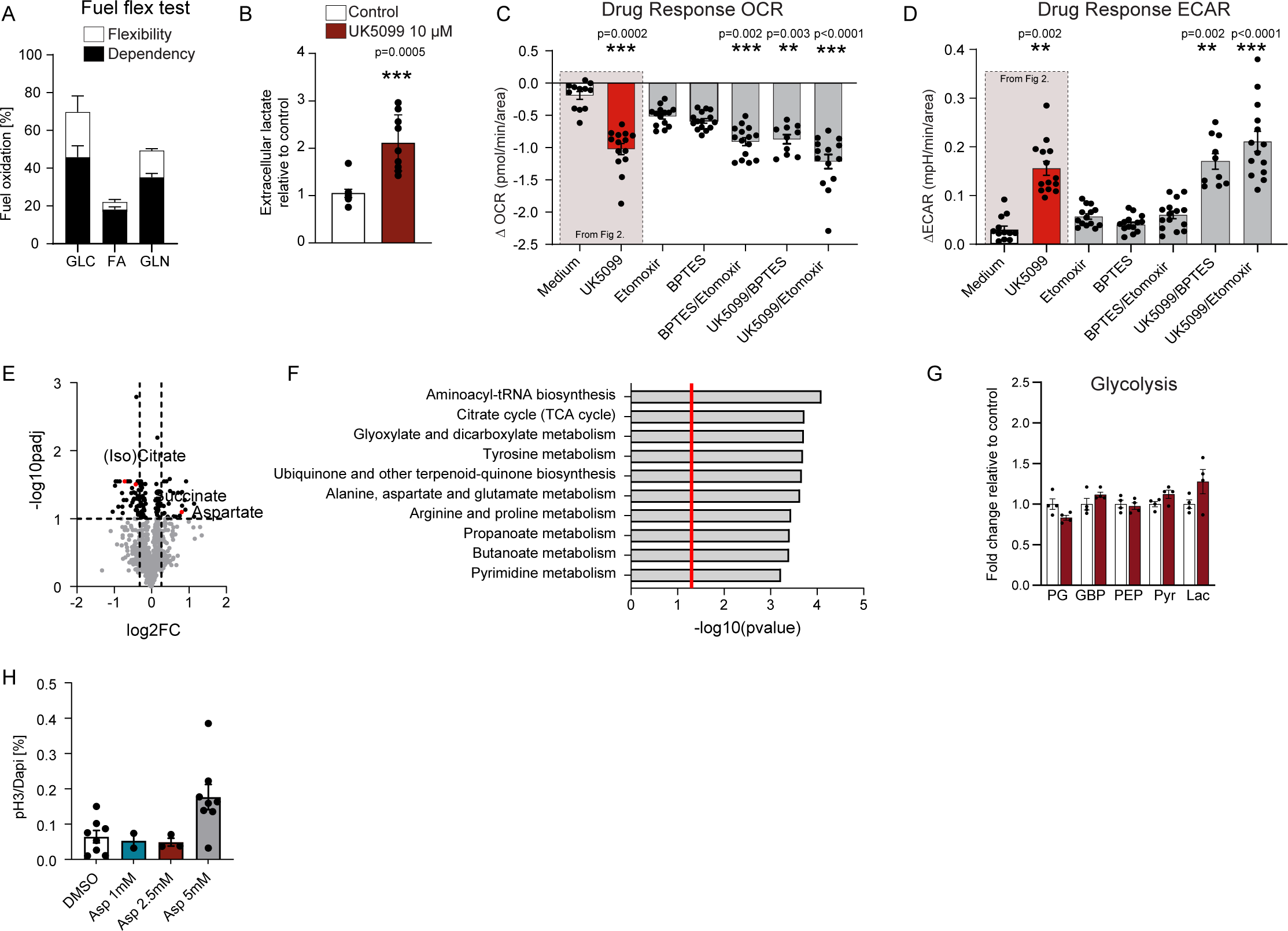
MPC1 inhibition activates quiescent NSPCs by increasing the intracellular pool of aspartate, related figure 2. A. Percent dependency (black) and flexibility (white) to oxidize glucose (GLC), glutamine (GLN) and fatty acids (FA) in quiescent NSPCs. Data represent mean ± SEM. n=3 independent experiments. B. Extracellular lactate concentration after 3 days of UK5099 treatment in quiescent NSPCs. Lactate concentration was normalized to protein content and normalized to control samples. Data represent mean ± SEM. ***p <0.001 n=9 samples from 3 independent experiments. Statistics were computed on averaged experiments (unpaired Student’s t-test). C. Difference in OCR after a single injection of medium (white), UK5099 1 μM (red), Etomoxir 4 μM, BPTES 3 μM, Etomoxir/BPTES, UK5099/BPTES or UK5099/Etomoxir in quiescent NSPCs. Data represent mean ± SEM. **p<0.01, ***p <0.001 n=10-15 wells from 3 independent experiments (one-way ANOVA with post-hoc test compared to medium, Statistics were computed on averaged experiments.) D. Difference in ECAR after a single injection of medium (white), UK5099 1 μM (red), Etomoxir 4 μM, BPTES 3 μM, Etomoxir/BPTES, UK5099/BPTES or UK5099/Etomoxir in quiescent NSPCs. Data represent mean ± SEM, **p<0.01, ***p <0.001. n=10-15 wells from 3 independent experiments (one-way ANOVA with post-hoc test compared to medium, Statistics were computed on averaged experiments) E. Representative Volcano plot of metabolomics data (fold change threshold = 1.2 and p-value adjusted=0.1) in control and UK5099 treated quiescent NSPCs. F. Pathway enrichment analysis of significantly changed metabolites between control and UK5099-treated quiescent NSPCs. Red line: p-value=0.05 G. Relative intensity of selected glycolytic metabolites in control and UK5099 treated quiescent NPSCs. Bars represent mean ± SEM. * adjusted p-value <0.1, N=4 biological replicates. PG: phosphoglycerate, GBP: Glyceric acid 1,3-biphosphate, PEP: phosphoenolpyruvate, Pyr: Pyruvate, Lac: Lactate H. Quantification of pH3 in quiescent NSPCs treated with several doses of aspartate. Data represent mean ± SEM. * p <0.05, n=2-8 CS from 3 independent experiments. (one-way ANOVA with post-hoc test compared to control.)

**Supplementary Figure 3.**
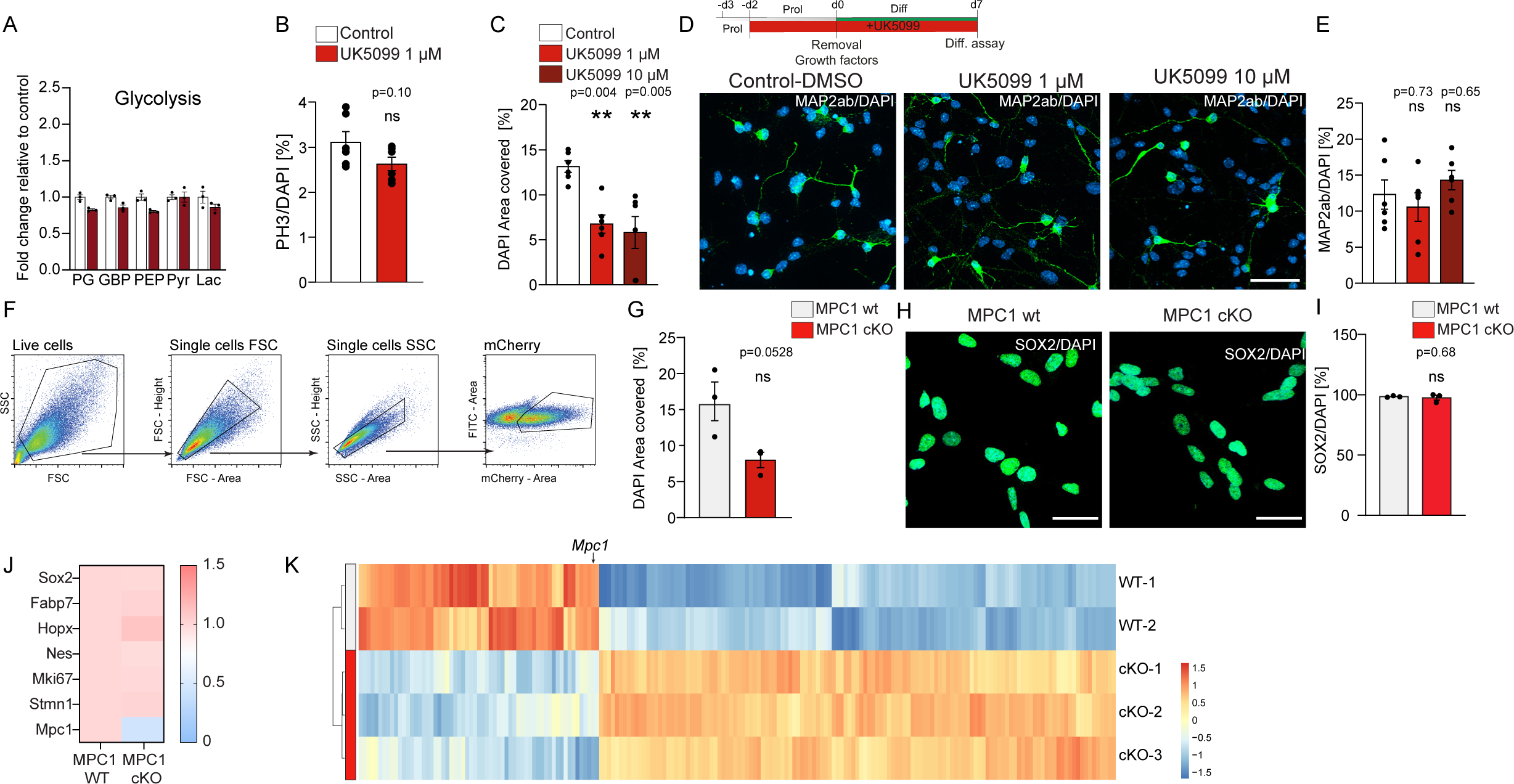
NSPCs lacking mitochondrial pyruvate important can still generate neurons through a shift in their metabolism, related to figure 3. A. Relative intensity of selected glycolytic metabolites in control and UK5099 treated proliferative NSPCs. Bars represent mean ± SEM. * adjusted p-value <0.1, N=4 biological replicates. PG: phosphoglycerate, GBP: Glyceric acid 1,3-biphosphate, PEP: phosphoenolpyruvate, Pyr: Pyruvate, Lac: Lactate B. Quantification of pH3+ in control (white) and UK5099-treated (red) proliferative NSPCs. Data represent mean ± SEM. Not significant (ns), n=6 CS from 2 independent experiments. (unpaired Student’s t-test). C. Quantification of DAPI area covered in control (white) and UK5099-treated (red) differentiated NSPCs. **p< 0.01. n=6 CS from 2 independent experiments. (one-way ANOVA followed by post-hoc test) D. Top: experimental outline of differentiation in presence of UK5099. Bottom: Representative confocal images of neurons (MAP2ab, green) and DAPI (blue) in UK5099-treated differentiated cells. Scale bar: 20 μm, E. Quantification of MAP2ab positive cells in control (grey) and UK5099-treated (red) cells. Not significant, n=6 CS from 2 independent experiments. (one-way ANOVA followed by post-hoc test) F. Sorting and gating strategy of MPC1 wt and MPC1 cKO mcherry positive cells. G. Quantification of DAPI area covered in MPC1 wt and MPC1 cKO cells. n=3 electroporations, mean ± SEM, unpaired Student’s t-test. H. Representative confocal images of SOX2 (green) and DAPI (blue) immunostaining in MPC1 wt and MPC1 cKO NSPCs. Scale bar: 50 µm I. Quantification of SOX2+ in MPC1 wt and MPC1 cKO NSPCs . Data represent mean ± SEM. Not significant (ns), n=3 electroporations (unpaired Student’s t-test). J. Heatmap of the fold change of the selected stemness markers and *Mpc1* in MPC1 wt and MPC1 cKO NSPCs. K. Heatmap showing the differentially expressed genes in MPC1-wt and MPC1-cKO NSPCs. arrow highlights *Mpc1* expression.

**Supplementary Figure 4:**
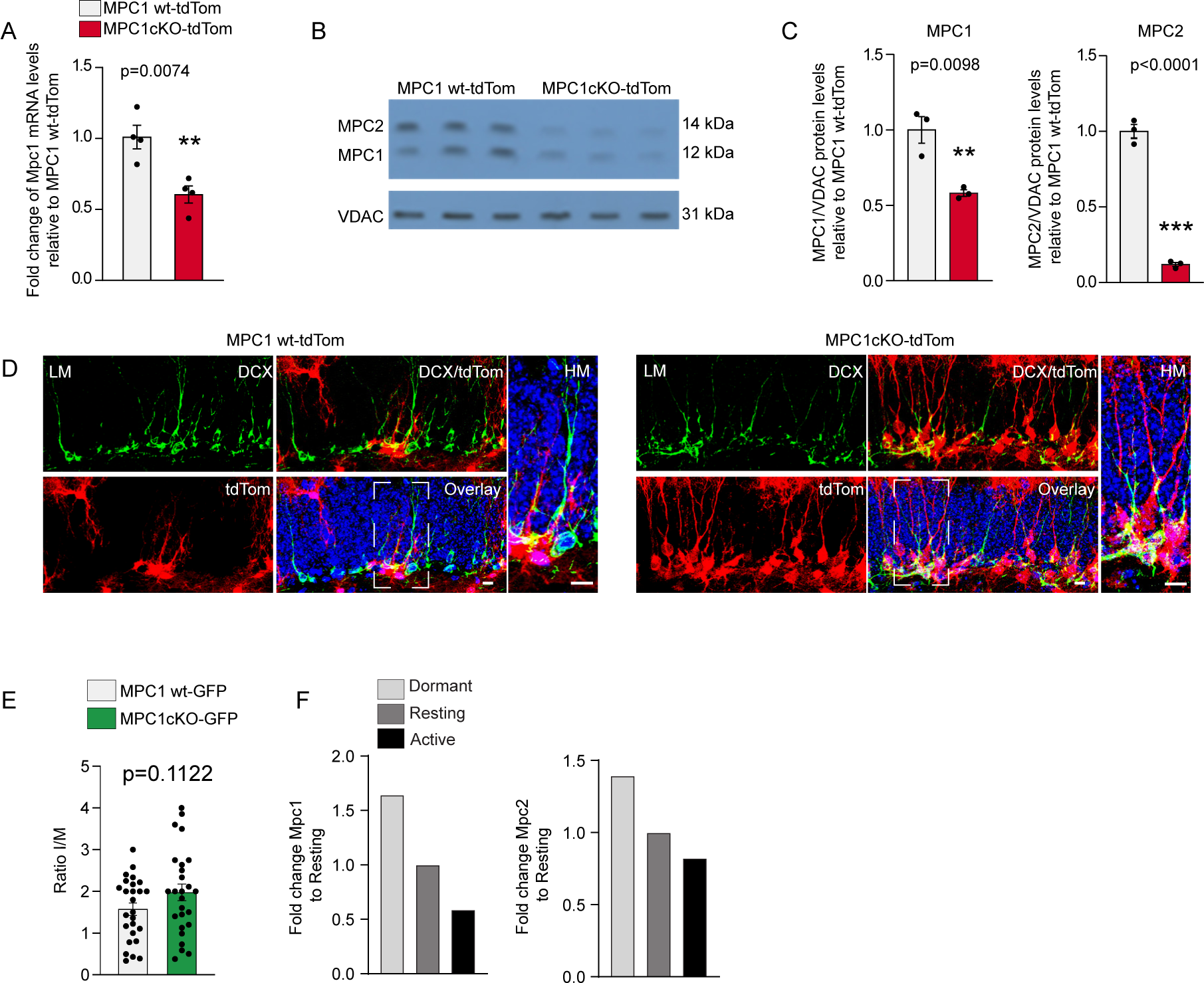
MPC1 deletion in NSPCs *in vivo* leads to increased numbers of progeny and increases neurogenesis, related to figure 4. A. Relative mRNA expression levels of Mpc1 from hippocampus of control MPC1 wt-tdTom (grey) and MPC1 cKO-tdTom (red) mice. Data are expressed as fold changed compared to MPC1 wt-tdTom and represented as means ± SEM. **p<0.01, n=4 mice per group (unpaired Student’s t-test). B. Representative western blots of MPC1 (12 kDa), MPC2 (14 kDa) and protein voltage-dependent anion channel (VDAC, 31 kDa) in the hippocampus of control MPC1 wt-tdTom (n=3 mice) and MPC1 cKO-tdTom mice (n=3 mice). C. Quantification of MPC1 and MPC2 protein expression in the hippocampus of control MPC1 wt-tdTom (grey) and MPC1 cKO-tdTOM mice. MPC1 and MPC2 expression was normalized to HSP70 levels and expressed as fold change to MPC1 wt-tdTom mice. Data are shown as mean value ± SEM. **p<0.01, ***p<0.001, n=3 mice per group (unpaired Student’s t-test). D. Representative LM and HM confocal images of TAM-induced td-Tomato recombination (red), Doublecortin (DCX, green) and DAPI (blue) immunostaining at P60 in the DG of control MPC1 wt-tdTom (grey) and MPC1 cKO-tdTom (red) mice. Scale bars: 10 μm. E. Ratio of the number of Immature/Mature (I/M) spines on Cre-induced tdTom+ newborn neurons in the DG of MPC1 wt-GFP and MPC1 cKO-GFP mice. Not significant (ns), n=4 mice per group, (unpaired Student’s t-test). F. Fold changes of *Mpc1* and *Mpc2* in dormant, resting and proliferating NSPCs. Processed scRNAseq data are from Harris et al. Cell Stem Cell 2021.

